# Continuous Serial Electron Diffraction for High Quality Protein Structures

**DOI:** 10.1101/2025.09.14.676192

**Authors:** Gerhard Hofer, Lei Wang, Laura Pacoste, Paul Hager, Alexis Fonjallaz, Lewis Williams, Emma Scaletti Hutchinson, Michele Di Palma, Pål Stenmark, Jonathan Worral, Roberto A Steiner, Hongyi Xu, Xiaodong Zou

**Author notes:** authors contributed equally (Lei Wang, Laura Pacoste).

## Abstract

Determining macromolecular structures is crucial for understanding biological mechanisms and advancing drug discovery. Three-dimensional electron diffraction (3D ED), also known as microcrystal electron diffraction (MicroED) using continuous sample rotation has emerged as a powful method for solving structures from sub-micrometre-sized crystals. However, the resolution of MicroED data from protein crystals is often limited by radiation damage. Serial electron diffraction (SerialED) overcomes this limitation by merging single-shot diffraction patterns from thousands of crystals, but its widespread use has been hindered by the complexity and scarcity of equipment required for single shot data acquisition. Here, we introduce continuous SerialED (c-SerialED) – a simple, robust and widely accessible protocol. This approach collects diffraction data quickly and efficiently from all crystals within a given area, without prior crystal identification. We show that only using a standard cryo-EM instrument equipped with a simple widely available CMOS detector, c-SerialED greatly reduces radiation damage while improving the data quality. We demonstrate that c-SerialED enables determination of lysozyme structures at atomic resolution (0.83 Å) and improves the data resolution of Dype Type Peroxidase Aa (DTPAa) crystals from 2.5 Å (MicroED) to 1.3 Å. Remarkably, the resulting structures are virtually free of radiation damage. The improved data quality and resolution allow visualization of radiation sensitive chemical features and protein-ligand interactions to state-of-the-art accuracy. By providing a convenient, fast, and damage-minimizing workflow on existing cryo-EM setups, c-SerialED significantly enhances the applicability of electron diffraction in structural biology. We anticipate our protocol will enable a wide range of studies requiring high-quality diffraction data from radiation-sensitive macromolecular crystals.

## Introduction

High-resolution, high-fidelity structural data are essential for reliably interpreting molecular function and guiding drug design. Over the past three decades, synchrotron-based X-ray crystallography has been the predominant technique for high-resolution structural analysis of macromolecules. Synchrotron facilities worldwide have implemented high-throughput workflows^1^, fragment-based screening for drug discovery^2^, serial crystallography^3^, and time-resolved crystallography^4^. However, obtaining macromolecular crystals with sufficient order and size for single-crystal X-ray diffraction (typically 5 × 5 × 5 μm^3^) remains a major challenge.

Electrons interact with matter much more strongly than X-rays,^5^ and can be readily focused using electromagnetic lenses, making them an ideal probe for both imaging and diffraction experiments. While recent advanced in cryo-EM single-particle analysis (SPA) have eliminated the need for crystallization^6,7^, the attainable resolution in most cases continues to lag behind that achieved by diffraction-based methods.

Three-dimensional electron diffraction (3D ED)^8^, also known as micro-crystal electron diffraction (MicroED), has been developed over the past decades as a powerful method for structural analysis of nano- and submicron-sized crystals that are too small for X-ray diffraction, which has revolutionized crystallography^9^. MicroED data collection is conceptually analogous to the rotation method in X-ray crystallography^1,2^: the crystal is continuously rotated in the transmission electron microscope (TEM) while diffraction patterns are recorded in movie mode on a fast detector.

Despite these advantages, major bottlenecks remain. The preparation and screening of high quality cryo-EM grids with protein crystals, as well as radiation damage caused by the electron beam^10^, still limit data quality and throughput. Specialized sample preparation techniques such as cryogenic focused-ion-beam (cryo-FIB) milling enable the preparation of high quality protein crystal lamella^11,12^, and the use of direct electron detectors allows collection of low-damage, high-resolution MicroED data^13^. However, these approaches require specialized, often costly instrumentation and are inherently low-throughput.

Serial electron diffraction (SerialED) is a relatively recent addition to the electron crystallography toolkit, with significant potential for the study of both inorganic crystals^14,15^ and macromolecular crystals^16^. In SerialED, single-shot diffraction patterns are collected from individual crystals distributed across the EM grid. The main advantage of SerialED is its ability to enhance the signal-to-noise ratio while drastically reducing electron beam-induced radiation damage. The main drawback, however, lies in sample preparation: EM grid must be densely populated with crystals of suitable size and morphology, which can be challenging to achieve reproducibly.

In current SerialED data collection protocols, a grid is first screened using TEM imaging^14^ or STEM imaging^15,16^. Crystals are then identified either manually or by automated imaging recognition algorithms. Electron diffraction patterns are then collected sequentially by combining electron beam and stage shifts to target the selected crystals. These methods require precise control of microscope optics and extensive calibration of crystal-recognition parameters, limiting both accessibility and throughput.

For macromolecular crystals embedded in vitrified ice, the inherent low contrast of cryo-EM micrographs further complicates the reliable identification of well diffracting crystal regions. Consequently, the need for rare nanocrystal samples, labour-intensive crystal targeting, and access of TEMs equipped for the required beam-control setup have collectively hindered the widespread adoption of this otherwise powerful technique.

Here, we introduce a continuous SerialED (c-SerialED) protocol for three-dimensional structural analysis of macromolecules that departs from the traditional paradigm of preselecting crystals prior to diffraction data collection. Instead, c-SerialED records diffraction data from entire regions of the grid, allowing crystals of interest to be selected retrospectively during data processing. In addition, we present practical methods for producing a dense slurry of suitably sized crystals, preparing grid manually from this slurry, and incorporating ligands when desired. Combined with our in-house software *Coseda* for automated data pre-processing and subsequent structure determination using established crystallographic software packages, this approach provides a complete workflow for high-quality protein diffraction experiments (Fig. 1).

**Figure 1.**
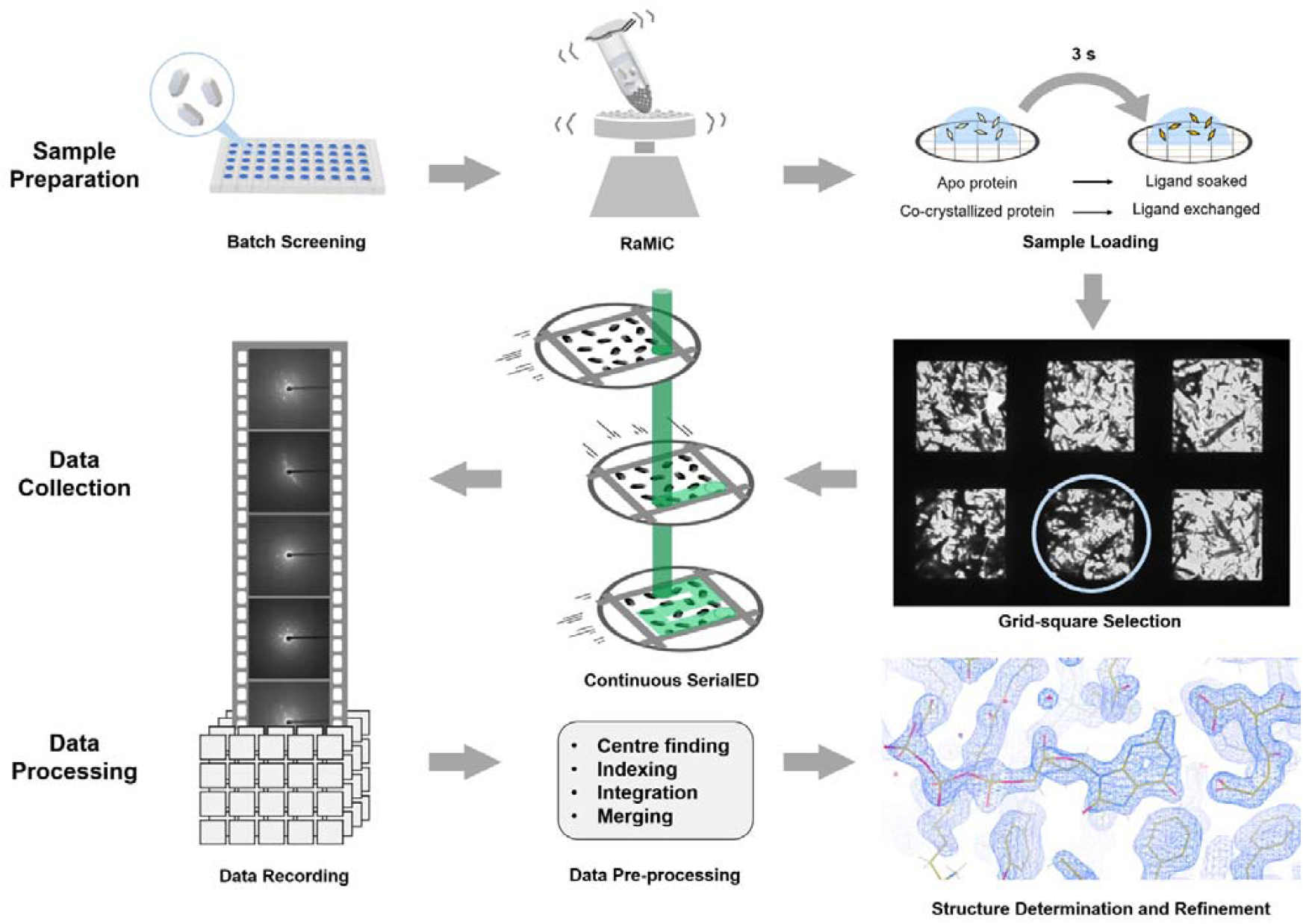
c-SerialED workflow. Batch-crystallised sub-micrometre crystals are deposited onto holey carbon EM grids. Through backside blotting, the crystals form a dense layer on the grid while excess mother liquor is removed. The grids are then plunge-frozen in liquid ethane, vitrifying the remaining solvent. During data collection, the frozen grid is navigated along a meandering rectilinear path by the microscope stage, while a parallel electron beam continuously illuminates crystals as they pass through the beam. Diffraction patterns are recorded in continuous acquisition mode on the detector. Frames containing diffraction signals are preprocessed and subsequently indexed and merged using CrystFEL, with final structure determination performed in PHENIX or CCP4.

### Sample preparation

Nanocrystals for c-SerialED were produced using our rapid-mixing protein crystallisation method (RaMiC), a modified batch crystallisation approach. In this protocol, repeated brief agitation with steel beads continually fractures growing crystals, thereby increasing the number of nucleation sites and restricting crystal growth to sub-micrometre thickness suitable for electron diffraction. The resulting crystal slurry is applied to cryo-EM grids using our detergent-assisted grid-preparation protocol. In this method, grids are first dipped in 1 % Tween-20 to render the carbon film hydrophilic and to promote uniform drainage, followed by manual plunge-freezing. Full experimental protocols are given in the Methods section.

### Continuous serial electron diffraction (c-SerialED)

For c-SerialED data collection, we raster-scan a selected region of the cryo-EM grid by continuously moving the stage in X/Y while recording electron diffraction patterns with a fixed, parallel nanobeam (300 nm to 500 nm in diameter). This procedure is conceptually similar to fixed-target serial crystallography at synchrotron beamlines^18^. In contrast to previous serial electron crystallography techniques, the beam is not shifted between exposures to target individual crystals. Instead, all frames are recorded as a continuous movie of the sample passing through the beam (Extended Data Fig. 1). This strategy maximises data yield by capturing all available diffraction signals and allows post facto selection of crystals of interest. It also lowers the technical barrier for implementation: whereas beam-shift-based methods require specialised hardware capable of correlated beam hopping and prior crystal identification (either manually or via automated software)^16^, our continuous-collection approach requires only stage control via a short script^19^ (see Supporting Information).

To address the problem of preferred crystal orientation on the support film, which can limit the sampling of reciprocal space, the grid can be tilted during data collection. Because c-SerialED uses a fixed beam position, crystals remain at eucentric height even when scanning across multiple grid squares. This allows the collection of complete datasets from samples exhibiting strong preferred orientation (see example below).

Our method collects all diffraction from a given area of the grid, regardless of crystal number or distribution. Consequently, a high crystal density is beneficial, as it increases the fraction of frames containing usable diffraction patterns. Using the grids prepared by RaMiC and our detergent-assisted protocol, we achieved hit rates between 5 % and 50 % in this study. Data collection throughput is primarily limited by detector speed, varying from 20 frames per second for scintillator-based CMOS detectors to several hundred frames per second for hybrid pixel detectors.

To generate MTZ files suitable for phasing in standard crystallographic software, the raw electron diffraction data are pre-processed using our in-house package *Coseda*. This pipeline performs peak finding, locates the direct beam position behind the beam stop, and corrects for beam drift during acquisition by using Friedel pairs. The resulting frame stacks are subsequently processed in CrystFEL^20^, where indexing is carried out with the XGANDALF^21^ algorithm, followed by integration and merging.

In summary, we developed a complete workflow that integrates rapid nanocrystal production using the RaMiC method, detergent-assisted cryo-EM grid preparation, c-SerialED data collection, and a data pre-processing pipeline *Coseda*. Structure determination is subsequently performed using conventional X-ray crystallography software, with indexing, integration, and merging in CrystFEL^19^, followed by phasing and refinement in PHENIX^22^ and the CCP4 package^23^.

### Complete High-resolution Data from Crystals with Preferred Orientation

The flat support film on which crystals rest often induces preferred orientation, limiting the sampling of reciprocal space when collecting diffraction from multiple crystals. This challenge is particularly pronounced for plate-shaped crystals and for crystals with low-symmetry space groups, which require extensive coverage of reciprocal space to achieve high completeness.

The triclinic form of hen egg white lysozyme (space group *P*1, a=26.66 Å, b=31.15 Å, c=33.57 Å, α=87.73°, β=108.97°, γ=111.6°) exemplifies this problem. To mitigate the effect of preferred orientation, we collected c-SerialED data at a static stage tilt of 45° (Fig. 2b). This tilt introduces additional variations in crystal orientation across the grid, while still relying on occasional crystal fragments with naturally deviating orientation to achieve complete data. Manual screening failed to locate crystals with rare non-aligned orientations. However, our continuous data-collection strategy imposes no selection bias, ensuring that any off-axis crystals are collected.

**Figure 2.**
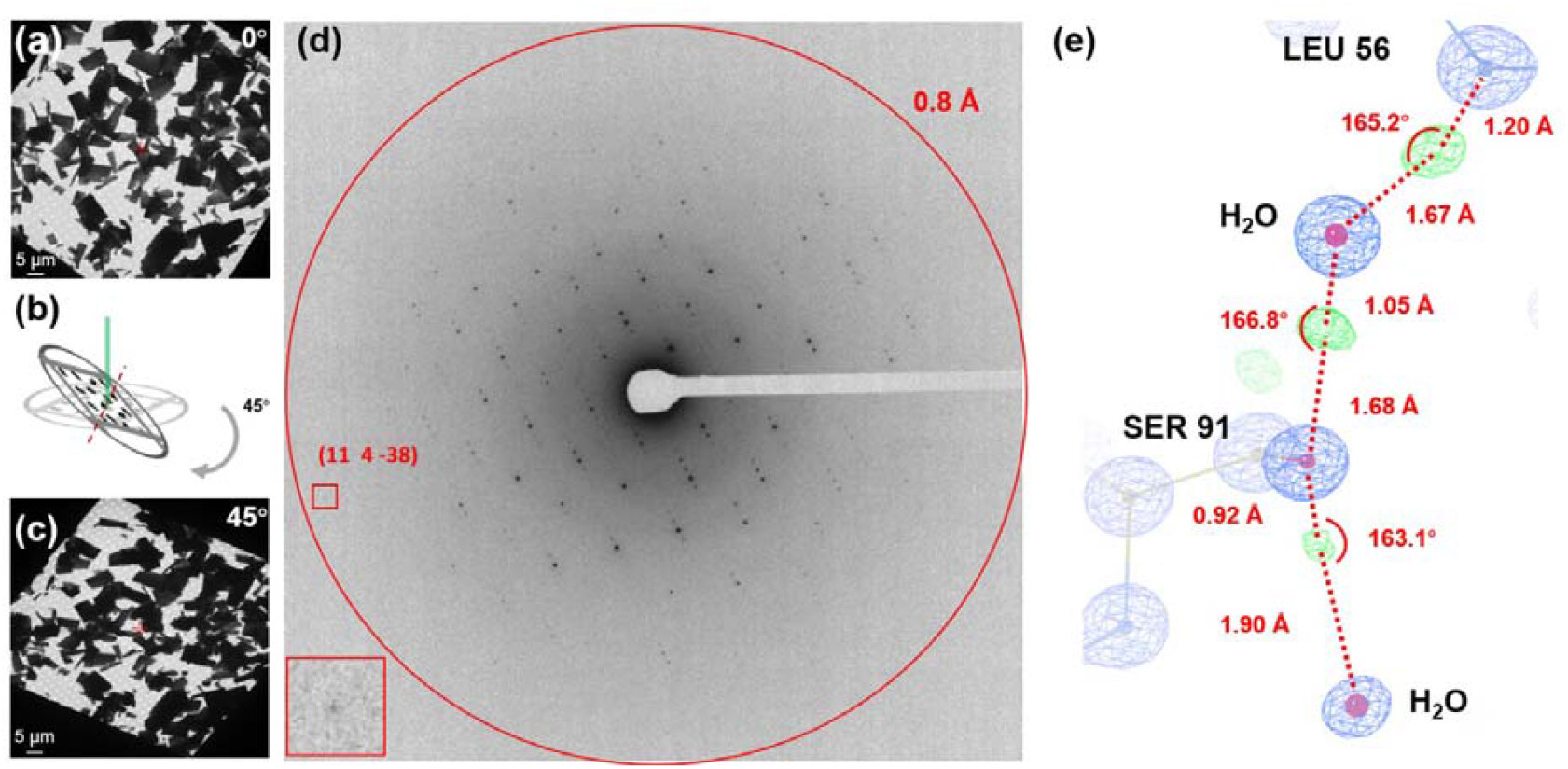
SerialED data acquisition of triclinic lysozyme using a CMOS detector. (a) Thin wedge-shaped lysozyme crystals on one grid square. (b) Schematic of the grid being tilted to 45° before data collection. (c) Grid square tilted to 45 °, under which tilt the SerialED data were acquired. (d) Sub-Ångström resolution diffraction pattern. A visible high-resolution peak can be identified in the red square. Insert: enlarged view of the spot. (e) Some H candidates for the solvent network can be identified from the difference map directly. The electrostatic potential 2F_o_-F_c_ map (blue) is contoured at 4 rmsd, while the F_o_-F_c_ difference map (green and red) is contoured at 3 rmsd.

To assess the highest achievable data quality for completeness, resolution and merging statistics using these triclinic plates at 45° tilt, we performed an extensive collection comprising 868,549 frames over a total of 24 hours without human intervention. As expected from the untargeted acquisition strategy, a substantial fraction of frames contain no diffraction, and additional frames were blocked by copper grid bars. Nonetheless, 38,122 patterns could be indexed, and after filtering, 30,794 frames were merged into the final dataset.

A representative high-resolution diffraction frame is shown in Fig. 2d. The merged dataset reached 99.0 % completeness at 0.83 Å resolution, which compares favourably with a recently reported 0.87 Å structure of this lysozyme form obtained by energy-filtered MicroED data collected on a direct electron detector ^13^. Using our data, we determined the structure of triclinic lysozyme at atomic resolution. The refined *2F*_*o*_*-F*_*c*_ map revealed well-resolved electrostatic potential for individual atoms. At this resolution, we anticipated to visualize hydrogen atoms, and indeed many hydrogens were observed, especially those forming the solvent network (see Fig. 2e and Fig. S3).

These results demonstrates that continuous data collection over all crystals on the grid at a fixed tilt of 45° enables the generation of nearly complete, high-quality electrostatic potential maps even for plate-shaped crystals in the lowest-symmetry space group. Although lysozyme crystals are relatively robust and diffract strongly, the combination of triclinic symmetry and preferred orientation represents a particularly challenging case, making this a stringent benchmark for our method.

### Radiation Damage Free Structures

Radiation damage has been a significant limitation in electron diffraction, causing loss of crystallinity and reducing the observable resolution after only a limited exposure. Site specific chemical changes can be observed before radiation damage leads to a visible loss of diffraction. This imposes a trade-off among completeness, signal-to-noise ratio, resolution, and radiation damage when collecting data from a single crystal. By concentrating all achievable diffraction for every crystal onto a single diffraction pattern, c-SerialED optimises the information extraction and should allow for the collection of less radiation damaged structures. To verify this and measure the visible damage, we applied the method to the study of Dye Type Peroxidase Aa from *Streptomyces lividans* (DtpAa)^24^ (*P*2_1_, a=72.77 Å, b= 67.40 Å, c=73.63 Å, α=90 °, β=105.8 °, γ=90 °).

DtpAa features a heme group with a water molecule completing the iron’s complexation shell. Previous studies have shown that the observed water-iron distance is sensitive to radiation-induced movement, with a linear trend between X-ray exposure and this iron-water distance^25^. RaMiC produced homogeneous crystal plates without the need for seeding. Data was collected both using MicroED and c-SerialED from dense grids prepared as described above.

By using c-SerialED, the achievable resolution improved from 2.5 Å for merged MicroED data of 20 crystals (out of 96 collected over 9 hours) to 1.3 Å for SerialED from 41,986 frames collected in 3 hours. Analysing the radiation-sensitive iron-water distance, the c-SerialED structure distance refines to 2.36 ± 0.07 Å, which is in good agreement with the 2.40 ± 0.13 Å distance observed in serial femtosecond crystallography (SFX), a method considered to be effectively free from radiation damage^26^. However, the MicroED structure distance is longer (3.29 ± 0.26 Å), indicating damage. Using the published correlation between this distance and radiation^25^, this movement corresponds to an X-ray exposure of 170-290 kGy (Fig. 3a). The drastic increase in achievable resolution observed from the *2F*_*o*_*-F*_*c*_ electrostatic potential maps (see Fig. 3 b-c), in addition to the absence of observable radiation damage, underscores the significant advantages of c-SerialED in attaining high-resolution and practically radiation damage-free structures.

**Figure 3.**
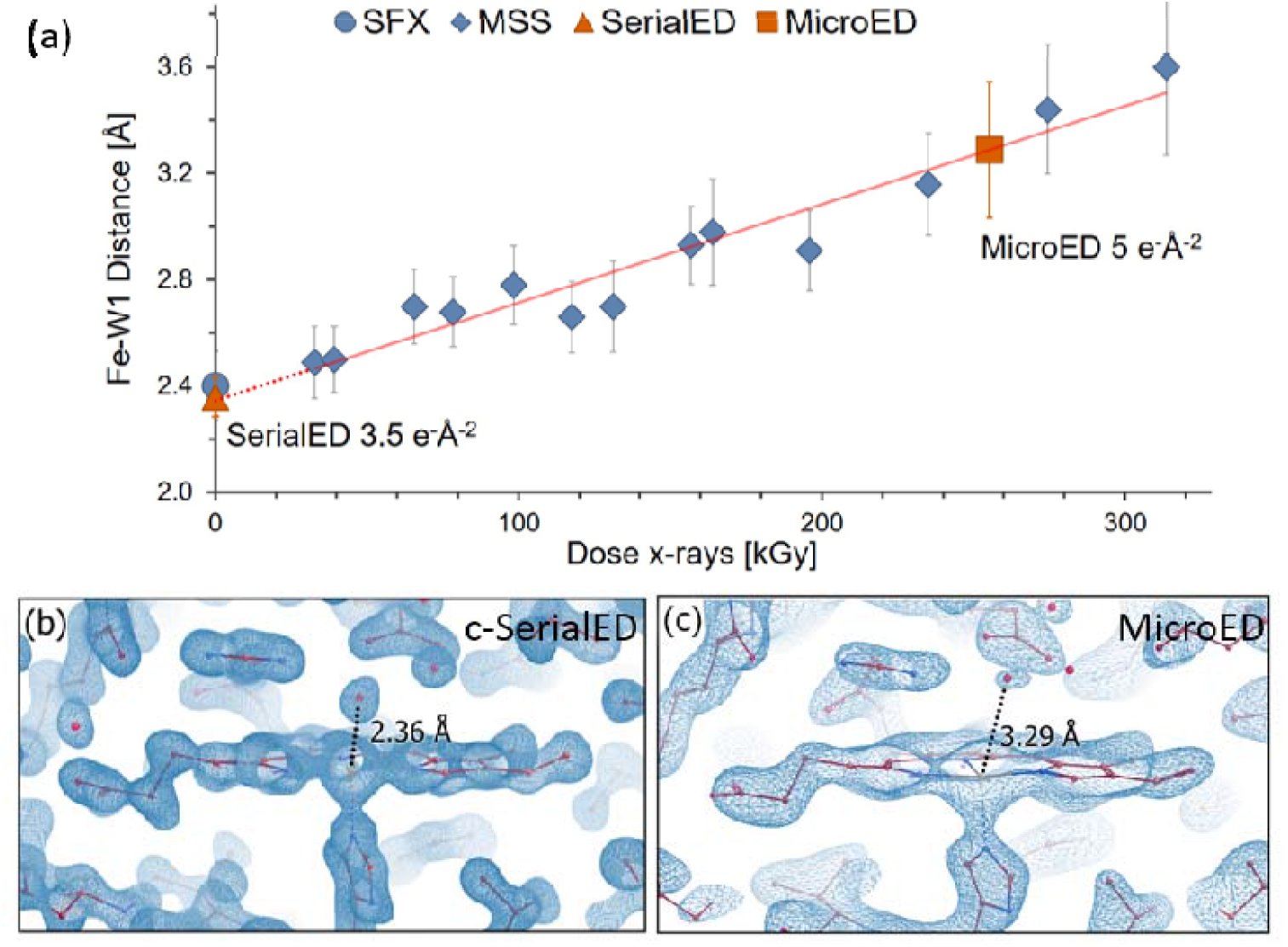
(a) Plot of Fe-H_2_O distance as a function of X-ray exposure from previously reported MSS series (gray diamonds) and SFX measurement (magenta diamond)^25^, fitted by a linear function (red line). Iron water distances with respective bond error are highlighted for c-SerialED (green horizontal line) and MicroED (red horizontal line) respectively. (b) 2F_o_-F_c_ maps of the c-SerialED (1.3 Å) and MicroED (2.5 Å) structures, non-filled and drawn at 1.5 rmsd, highlighting the shorter water-iron distance in the c-SerialED structure.

### On-Grid Ligand Soaking and Exchange

Ligand soaking is a valuable tool for drug discovery, allowing protein-ligand interactions to be probed at atomic resolutions. However, they can be time-consuming as large crystals can be destroyed by the stress of non-uniform binding caused by steep diffusion gradients^27,28^. The thin crystals used in c-SerialED are far more resilient to this stress due to the rapid diffusion on the sub-μm scale. In enzyme/substrate soaking, this fast diffusion also avoids differing turnover states throughout the thickness of the crystals. Additionally, slight mismatches in condition (pH, salinity, water activity) can cause a slow dissolution of the crystals during longer soaking, but can often be withstood for the short contact time (< 5 seconds) of on-grid soaking.

Once prepared, the crystal slurry produced by RaMiC allows for rapid introduction of new ligands to pre-grown protein crystals on the grid, streamlining ligand binding experiments^29^.

For instance, when sodium azide is introduced to DtpAa crystals for 5 seconds before freezing, the resultant electron density map distinctly reveals the azide ion substituting for the water molecule at the heme binding site (Fig. 4a and b). Our manual grid preparation not only allows for swift ligand soaking, as is commonly used for single crystal X-ray diffraction studies, it also enables the complete exchange of the original crystallisation ligand for a new ligand with remarkable ease.

**Figure 4.**
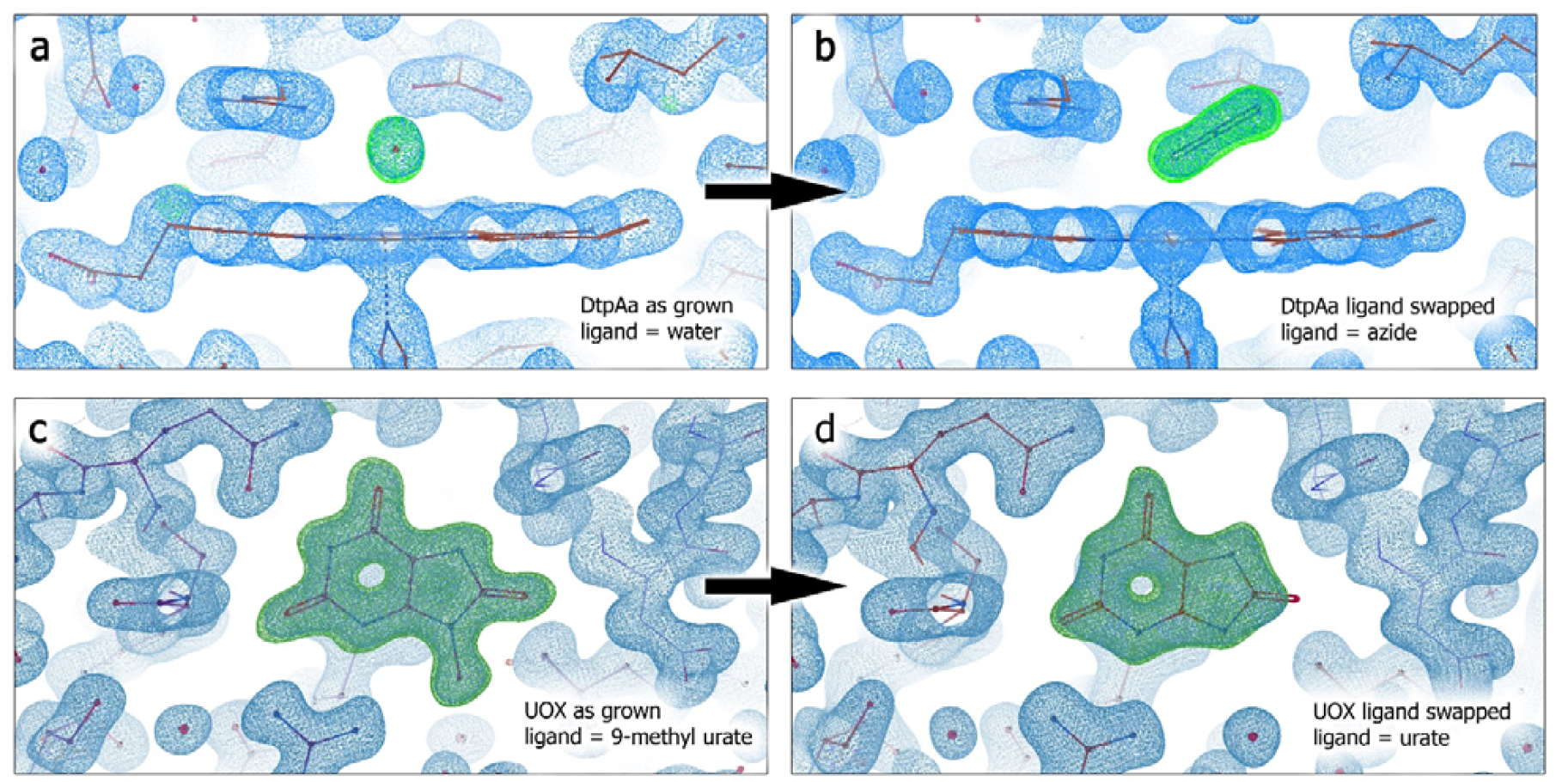
On-grid ligand exchange visualised by continuous SerialED omit maps (a) DtpAa as crystallised with bound water omitted. (b) DtpAa on grid water to azide swap omit map. (c) UOX as crystallised with 9-methyl urate omit map. (d) UOX on grid ligand swap to urate omit map. All 2F_o_-F_c_ maps are non-filled and drawn at 1.5 rmsd (blue), F_o_-F_c_ maps at 4 rmsd (green,red)

For on-grid ligand swapping, after the excess liquid is removed by the first back-side blotting, 2 μl of precipitant solution with the desired new ligand is pipetted onto the grid. After optionally waiting for three seconds, the grid is blotted for a second time before being vitrified as discussed above. In the case of urate oxidase (UOX)^30,31^, which requires a ligand to occupy the active site to form the well diffracting *1*222 polymorph (*1*222, a=80.58 Å, b= 94.49 Å, c=103.89 Å, α=90 °, β=90 °, γ=90 °). Using its substrate of uric acid would result in degradation products in the final structure (unless crystallised under anaerobic conditions). However, a ligand like 9-methyl uric acid (9-MUA) can be used during the crystallisation (Fig. 4c)^32^. These crystals can then be flushed with uric acid once deposited on the grid. This procedure results in a thorough replacement of 9-MUA with the substrate at the active site, as clearly demonstrated by the absence of the methyl group seen in Fig. 4d compared to Fig. 4c.

The collection strategy used for the 9-MUA bound UOX structure aimed at providing high resolution and map quality through high redundancy in observation. 898,649 frames were collected of which 56,538 were merged in a complete set of data and a structure was refined to 1.25 Å. For the urate swapped structure of UOX only 3,823 diffraction patterns were collected over roughly 45 minutes, showing that while lower redundancy provides lower completeness and resolution, ligand binding clarity can be obtained in a short amount of time. The final dataset of 96.5 % completeness was refined to 1.75 Å.

The ability to quickly and efficiently swap ligands as needed, coupled with the uncomplicated data collection of c-SerialED opens the door for fast fragment screening as well as time-resolved measurements of slow enzymatic reactions in a highly automatable way.

### Protein-Ligand Complex of Cancer Drug Target

Accurate substrate-binding geometries provide the essential blueprint that underlies nearly every structure-based drug-discovery and screening strategy. To provide an example for the application of the complete c-SerialED procedure outlined above and to compare it to MicroED using the same microscope and crystals, we analysed the binding of the natural substrate 8-Oxo-2’-deoxyguanosine-triphosphate (8-oxo-dGTP) to MutT homologue 1 (MTH1).

This project required data collection from flat crystals with preferred orientation, soaking of the substrate into apo crystals to prevent enzymatic degradation, avoiding radiation damage, and pushing the resolution past 2 Å to clearly visualize the binding mode of the ligand.

MTH1, a Nudix-family enzyme, hydrolyses mutagenic nucleotides, maintaining genomic integrity and playing a key role in cancer biology and aging, making it a valuable structural biology target^33–35^.

Apo crystals of MTH1 (*P*2_1_2_1_2_1_, a= 59.34 Å, b= 67.55 Å, c= 80.11 Å, α=90 °, β=90 °, γ=90 °) were grown using RaMiC. Immediately before plunge freezing, the substrate 8-oxo-dGTP was soaked into crystals on-grid for 3 seconds to prevent enzymatic hydrolysis. In our previous structural studies of MTH1, we were only able to obtain crystals large/thick enough to collect good quality X-ray diffraction data when we co-crystallized the enzyme in the presence of substrates/ligands^33–37^. Importantly, the apo MTH1 crystals used for c-SerialED here, which were not suitable for X-ray crystallography data collection, generated structural data of excellent quality. Additionally, as it can take up to a week for crystals large enough for X-ray crystallography to grow, the risk of substrate hydrolysis increases, and often only the monophosphate product is observed in the electron density^33–35^.

c-SerialED on grids prepared with our detergent-assisted protocol involved 4 hours of data collection at 45° tilt. Processing over 1 day resulted in a 1.66 Å resolution structure (Fig. 5a). The MicroED data collection with a high-dose (total fluence of up to 7.4 e^-^Å^-2^ per crystal) on the same grid, chosen to provide the highest resolution possible, provided a map within only 2 hours to 2.32 Å.

**Figure 5.**
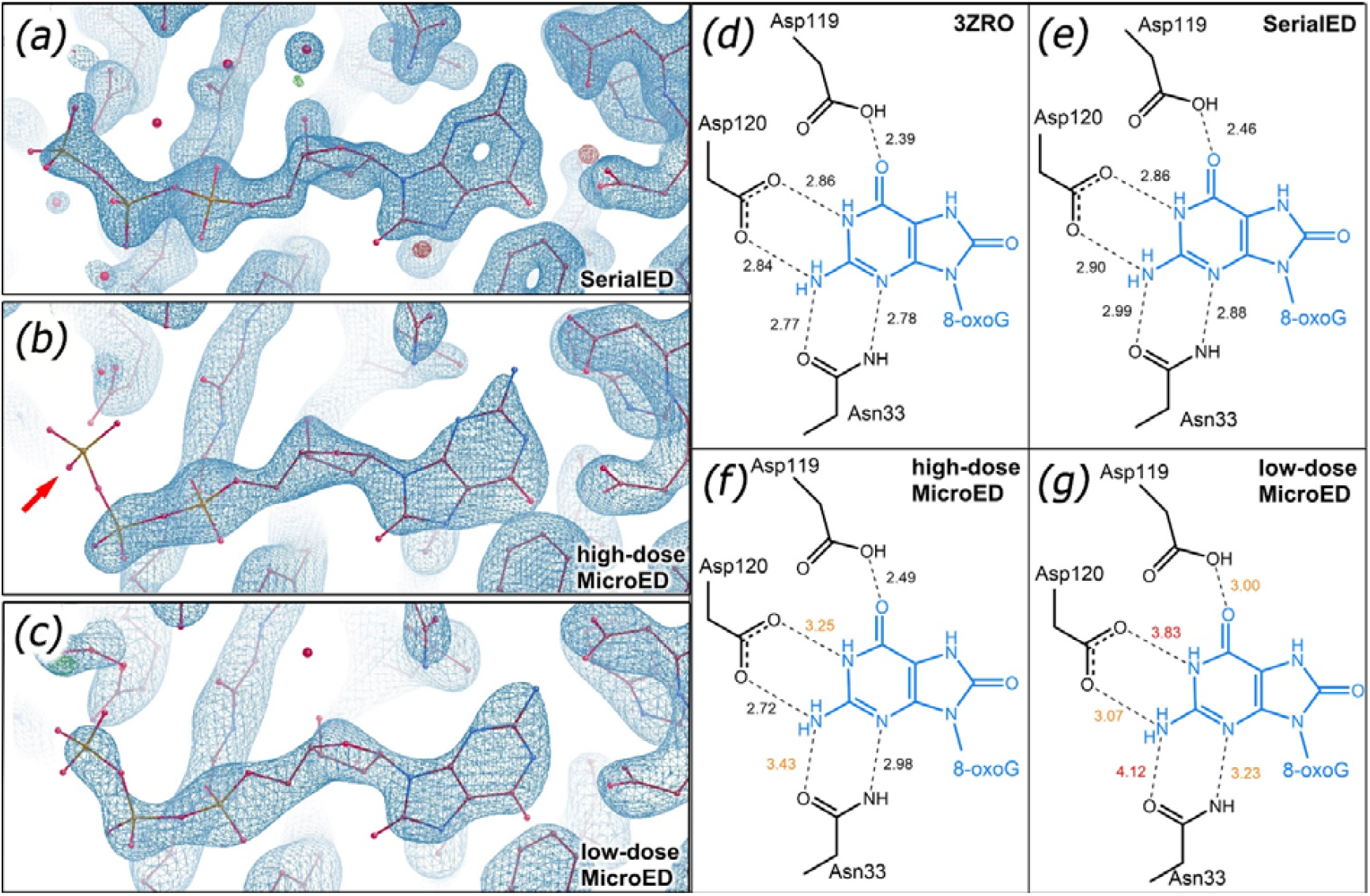
MTH1 maps resutling from (a) c-SerialED refined to 1.66 Å, (b) high-dose MicroED refined to 2.32 Å with the γ⍰ phosphate pointed out by a red arrow, and (c) low-dose MicroED refined to 2.86 Å. All 2F_o_-F_c_ maps are non-filled at drawn at 1.5 rmsd (blue), F_o_-F_c_ maps at 4 rmsd (green,red). (d) Distances between hydrogen bonding partners of the previously published X-ray structure (PDB code 3ZR0) compared to their respective distances in the three electron diffraction structures. Distances are marked in black if under 3.0 Å,

Comparing the two structures showed a clear lack of signal of the γ-phosphate in the MicroED structure (Fig. 5b), presumably due to radiation damage. Collecting an additional set of MicroED data at a lower dose, keeping the fluence below 1.1 e^-^Å^-2^ per crystal, restored the signal at the terminal phosphate group, yet also dropped the usable resolution to 2.86 Å structure (Fig. 5c). This showcases the trade-off between signal strength providing higher resolution and the damage incurred by the stronger illumination using the MicroED method.

The binding mode of 8-oxo-dGTP to MTH1 is visualised in Fig. 5d. In addition to a pi-stacking interaction with Trp117, the base of 8-oxo-dGTP is supported by hydrogen bond interactions with Asn33, Asp119 and Asp120. These two aspartate residues are important for the broad substrate recognition of MTH1, as they can change protonation state depending on whether a guanosine or adenine base is bound^38,39^.

When comparing hydrogen bond distances between key residues (Asn33, Asp119, Asp120) and 8-oxo-dGTP, our c-SerialED structure showed the highest similarity to a 1.80 Å X-ray structure (PDB code 3ZR0) in Fig. 5 d and e. The MicroED structures, particularly the low-dose structure, showed notably less accurate distances as shown in Fig. 5 f and g. This is primarily due to the drastically lower resolution not allowing for an atomically correct placement of the ligand and interacting side chains. The high-dose MicroED structure regains some of the positional accuracy due to the higher achieved resolution but shows clear signs of radiation damage. Only the c-SerialED structure was of sufficient quality to unambiguously determine the substrate binding mode.

## Discussion and Outlook

Our c-SerialED method addresses several long-standing bottlenecks in electron diffraction (MicroED and SerialED) data collection. It eliminates the need for extensive screening for high-quality crystals on the cryo-EM grid, mitigates radiation damage, and reduces the impact of preferred crystal orientation. By employing a high-throughput raster-scan strategy, c-SerialED efficiently collects diffraction data from all crystalline material within a selected region of interest, without introducing selection bias.

Because the number of collected frames scales linearly with acquisition time, users can tailor data collection to the scientific question at hand. If the goal is to obtain the best achievable structure, c-SerialED can be run for extended collection periods to maximise crystal coverage. Data from the best diffracting crystals, including those in rare orientations, can then be merged to yield the highest possible structure quality. For example, we merged 33,745 patterns for the lysozyme dataset (Fig. 2e), achieving high completeness at atomic resolution. Similarly, for the 9-MUA bound UOX structure, 898,649 frames were collected, of which 56,538 were merged, producing a highly redundant dataset that refined to 1.25 Å resolution (Fig. 4c).

On the other hand, when rapid structure determination is desired, a smaller region can be selected, reducing both data collection and process time. For the urate-swapped UOX structure, only 3,823 diffraction patterns were merged from data acquired in 45 minutes. Although the final dataset had slightly lower redundancy and completeness (96.5 %) and refined to 1.75 Å, the ligand density was sill clearly interpretable (Fig. 4d), demonstrating that that c-SerialED can provide biologically meaningful results even under time constraints.

Importantly, the simplicity of our data-collection strategy, combined with our data processing pipeline, makes c-SerialED broadly accessible to laboratories equipped with a TEM, basic automated stage control, and a diffraction-capable detector. While the method requires a sufficiently dense populations of nanocrystals, it offers superior achievable resolution compared with MicroED and minimises radiation-damage artefacts associated with rotation-based data collection. Notably, despite the overlapping illumination areas between consecutive frames inherent to continuous data collection, we observed no detectable signs of radiation damage under our acquisition conditions (Fig. 3).

Although protein-ligand complexes have previously been solved by MicroED^40,41^, our approach advances ligand screening by combining straightforward ligand incorporation during manual grid preparation with high-quality data. The resulting omit maps (Fig. 4) demonstrate accurate ligand placement, enabling precise interpretation of substrate and product binding modes, which is critical for elucidating enzyme reaction mechanisms. Moreover, because c-SerialED introduces significantly less radiation damage than both MicroED and X-ray crystallography, it may be particularly well-suited for proteins containing metal ions or redox-active cofactors that are radiation sensitive.

These features make c-SerialED attractive for structure-based drug design. Although it is currently less high-throughput than synchrotron-based X-ray crystallography, it could be readily combined with complementary fragment-screening approaches such as differential scanning fluorimetry (DSF), enzymatic assays, or surface plasmon resonance (SPR) to identify hits, whose binding modes can then be solved using c-SerialED. The short soaking times required are also advantageous for crystals that are incompatible with prolonged exposure to DMSO-containing solutions, which are common in fragment libraries.

While the present work focuses on macromolecular crystallography, the approach is equally applicable to small-molecule samples, and we are actively developing this capability. A user-friendly graphical interface for our processing pipeline is in progress to further lower the barrier to adoption. Taken together, the simplicity of the c-SerialED collection scheme, its minimal instrumentation requirements, and the reproducible RaMiC sample-preparation protocol have the potential to democratize high-quality electron diffraction. We anticipate that this method will enable laboratories worldwide to obtain high-resolution maps more efficiently, accelerating discoveries in structural biology and chemistry.

## Acknowledgements

This work was supported by the European Union’s Horizon 2020 research and innovation programme under the Marie Sklodowska-Curie grant agreement no. 956099 (NanED − Electron Nanocrystallography−H2020-MSCAITN). We also acknowledge financial support from the Swedish Research Council (VR, 2019-00815 (XZ) and 2022-03596 (HY)), and the Knut and Alice Wallenberg Foundation (KAW, 2019.0124). The c-SerialED and MicroED data were collected at the Cryo-EM Swedish National Facility funded by KAW, Family Erling Persson and Kempe Foundations, SciLifeLab, Stockholm University and Umeå University. We also thank Dr. Mathieu Coinçon, Dr. Dustin Morado and Dr. Marta Carroni for TEM support at Scilifelab. MDP is a PhD student funded by the Italian Ministero dell’Università e della Ricerca (MUR).

