## Supporting Information for "Continuous Serial Electron Diffraction for High Quality Protein Structures"

† authors contributed equally

* corresponding authors

### Rapid Mixing Crystallization (RaMiC)

In SerialED, each piece of crystal is only diffracted once. It is therefore necessary to start out with a large number of crystals of appropriate morphology on a cryo-EM grid. We developed an unconventional approach of agitated batch crystallization whereby an exponentially growing number of seed crystals limits the crystal growth. The approach is similar to previously published protocol for growing microcrystals suitable for X-ray free electron lasers (XFEL)[1], but optimized to produce smaller crystals (<500 nm thickness) specifically suited for electron diffraction. Our approach, named Rapid Mixing Crystallization (RaMiC), involves repeated rounds of agitated mixing with steel beads during crystal growth, to maintain crystals within the desired size limit.

Full details on the development of the RaMiC procedure are to be published. In short, initial crystallization conditions are identified by conventional protein crystallization screening in 1 µL drops before moving to larger batch crystallization to save on the total protein required. These conditions are then recreated in 200 µL tubes with the addition of roughly 20 steel beads of 0.5 mm diameter. These are used to both agitate the crystal suspension immediately after the addition of the protein to the precipitant mix, as well as to create additional seeds from crystals growing during this experiment. Adding seeds to the batch setup can be helpful but is not required in all cases. After the addition of the protein solution, the tube is touched to an orbital shaker for 5 seconds to quickly reach uniform mixing. This mixing is repeated every few minutes as long as crystals are growing. Since larger crystals are more likely to be damaged by the shear forces of this procedure and yield additional seeds, this increases the number of seeds exponentially with every step until the crystal growth is stopped by the end of protein super-saturation. The exact crystallization conditions, including whether seeds are used, vary between samples, and full details for each case are provided in the section 'Sample Preparation Details'.

If successful, the final product of the RaMiC procedure is an opalescent suspension of microcrystals, This suspension is directly used for diffraction data collection, instead of just providing seeds for final batch crystallization as is the case in other protocols[1]. The crystal slurry produced by RaMiC can be spun down using a PCR-tube centrifuge, to sediment any large crystal pieces, before applying up to 2 µL to a cryo-EM grid with holey carbon foil.

### Detergent-Assisted Grid Preparation

For grid preparation of protein microcrystals, the hydrophobic TEM grid is typically made hydrophilic using a low-energy plasma, for instance by using a glow discharge, to ensure an even distribution of the aqueous crystal suspension.[2] Alternative approaches to glow discharge, such as coating the grid with a surfactant to fixate hydrophilic samples[3], have been used to prepare samples for TEM imaging. However, to our knowledge, this has never been applied to the preparation of protein microcrystals for electron diffraction data collection. We found that dipping the grid in a solution of a mild non-ionic detergent (e.g. 1 % Tween-20) prior to sample deposition was sufficient to make the carbon film hydrophilic, eliminating the need for glow discharging. At the same time, dipping results in coating of both sides of the grid, making the backside hydrophilic as well, which improves drainage of the mother liquor during blotting.

After applying the crystal suspension to the grid, the backside of the grid is touched to a filter paper, allowing excess mother liquor to pass through the holey carbon foil. A layer of dense crystals covered by a thin film of liquid is left on the grid which is manually plunged into liquid ethane for vitrification. This procedure requires no machines, works equally well for PEG and high salt conditions, and takes less than a minute per grid.

### Sample Preparation Details

#### 1. Lysozyme

Lysozyme powder (Sigma-Aldrich) was dissolved in 0.05 M, pH 4.5 sodium acetate buffer. Two lysozyme solutions were prepared at concentrations of 20 mg/mL and 40 mg/mL, respectively. Sodium nitrate was also dissolved in the same buffer to achieve concentrations of 0.4 M and 0.8 M.

(a) Initial seed preparation was based on the protocol described in the previous study[4]. 20 µL lysozyme solution (20 mg/mL) and 20 µL sodium nitrate solution (0.4 M) was added to a PCR tube, without mixing. The tube was then placed in a refrigerator at 4°C overnight. After this incubation period, the tube could be stored at room temperature. Within 3-5 days, several large, irregular cubic crystals formed. To prepare suitable seeds, 0.5 mm steel beads were added to the tube, and the large crystals were fragmented into smaller pieces using a vortex machine. Agitation was continued for several tens of seconds until the solution became opaque.

(b) Nano-sized crystallization was initiated by directly pipetting 0.5 µL of seed slurry into a mixture of 20 µL lysozyme (40 mg/mL) and 20 µL sodium nitrate (0.8 M) in a PCR tube containing 0.5 mm steel beads. The tube was then immediately touched by a vortex machine. Mixing was performed for 30 seconds per round until the solution became milky. Before loading the sample onto a TEM grid, crystal slurry was spun down to sediment oversized crystals.

TEM grids were first prepared by dipping them into a 1% Tween-20 solution, followed by removing the excess detergent through back blotting. Subsequently, 5 µL crystal slurry was loaded onto the TEM grid and backside blotted. As soon as the liquid was drawn through the grid, 2 µL of 2.25 M NaCl in 100 mM sodium acetate pH 4.5 was pipetted onto the grid as cryoprotectant. The grid was then blotted from the backside again for 3-5 seconds and plunge-frozen in liquid ethane manually.

**2. DtpAa**

For crystallization, 10 µL 25 % PEG 3350 100 mM HEPES pH 7 were pipetted into 30µL of 31 mg/mL DtpAa Y398F in a PCR tube without mixing. No seeds were added to the mixture. After 3 minutes, in which a cloudy interface layer forms between the two liquids, the tube is touched to a vortex mixer for 10 seconds. The resulting crystal slurry was of very uniform crystal size and did not require centrifugation before application onto a TEM grid. 1 µL of the slurry were applied to a Quantifoil QF 0.6/1 300 Cu grid treated by dipping in 1% Tween-20 and blotting off excess liquid, before manually backside blotting the mother liquor for about 3 s before manually plunge freezing in liquid ethane. For the azide bound crystals, 2 µL of 100 mM sodium azide in 10 % PEG 3350 100 mM HEPES pH 7 were pipetted onto the crystals on the grid after blotting and the blotting repeated before plunge freezing.

**3. MTH1**

**Crystallization**

For crystallization, 0.5 µL protein solution (14 mg/mL) was first mixed with 0.5 µL of precipitant solution (30% PEG-6000, 0.16 M $Li_2SO_4$, 0.1 M sodium acetate, pH 4) in a batch plate. Needle-like crystals appeared within 1 hour. To prepare seed crystals, 1 µL of these crystals was transferred to 15 µL of precipitant solution and ground using steel balls. Subsequently, 15 µL of protein solution was added to the prepared seed solution and vortexed for 5 s to generate microcrystals.

**Ligand Soaking on the TEM Grid**

To prepare the ligand solution, a precipitant solution diluted to 50% of its original concentration (15% PEG-6000, 0.08 M $Li_2SO_4$, 50 mM sodium acetate, pH 4) was used. 8-Oxo-dGTP and $MgCl_2$ were then added to reach final concentrations of 1.5 mM and 3 mM, respectively. Ligand soaking was performed on a C-flat 1.2/1.3 200-mesh TEM grid. The grid was pre-treated by dipping into a 1% Tween-20 solution. 1 µL of dense microcrystals was diluted in a mixture of 5 µL precipitant solution and 5 µL Milli-Q water. 1.5 µL of diluted crystal slurry was then applied to the TEM grid. The grid was back blotted with filter paper to remove excess liquid. 1.5 µL of ligand solution was immediately added to the grid. After a 3-second ligand soaking, excess ligand solution was removed by back blotting, and the grid was plunge-frozen in liquid ethane.

**4. UOX**

For RaMiC, 15 µL protein solution (22 mg/mL) saturated with 9-methyl-uric acid (2 h incubation with excess powder) was mixed with 0.5 µL of seed solution and 15 µL precipitant solution (30 % PEG 3350, 100mM Tris Acetate pH 8, 100mM $Na_2SO_4$) PCR tube containing 0.5 mm steel beads. The tube was then immediately touched by a vortex machine for 10 seconds. After 5 minutes 6 µL of 50% PEG 3350 were added and the mixing repeated. The resulting slurry was left over night before use. 1 µL of the top layer after 15 minutes of settlement time were transferred onto C-flat 1/1 200-mesh TEM grid dipped in 1% Tween-20 solution. For the methylurate dataset, the grid was backside blotted for approximately 3 s until the solution had visibly passed through the grid. For the urate data set 2 µL of 20 % PEG 3350, 100mM Tris Acetate pH 8, 100mM $Na_2SO_4$ saturated with uric acid were added at this stage and the backside blotting repeated. This urate wash was repeated a second time before the grid was manually plunged into liquid ethane.

**Instrument**

c-SerialED and MicroED data were acquired using a Titan G3i transmission electron microscope operating at 300 keV. A Ceta-D detector was used for data collection.

**c-SerialED Data Collection**

c-SerialED data was collected with settings as presented in Extended Data Table 1**.** During data collection, the beam was held constant while the stage was moved to scan the grid, with or without prior tilt of the grid (Extended Data Figure 1). Diffraction patterns were recorded simultaneously using *Velox* for camera control.

Stage sweeping was performed using a 5-line script in *SerialEM*[5]. The main lines of the script followed the general structure:

*"MoveStageWithSpeed [distance in µm in x-direction] [distance in µm in y-direction] [speed factor]"*

One example of our 5-line script is shown below:

*"MoveStageWithSpeed 40 0 0.0625*

*MoveStageWithSpeed 0 1 1*

*MoveStageWithSpeed -40 0 0.0625*

*MoveStageWithSpeed 0 1 1*

*Repeat"*

In this script, the first line moves the stage 40 µm in the x-direction, with no movement in the y-direction, using a speed factor of 0.0625. The speed factor can be set within a range of 0 to 1. In our setup, a factor of 0.0625 corresponded to a speed of approximately 3.73 µm/s. The speed was adjusted depending on the beam size and the exposure time per frame, which varied depending on the sample (Extended Data Table 1). For the data presented here, the speed factor was chosen to ensure a movement of half the beam diameter per frame, generating overlap of the sample area recorded on each frame (Extended Data Figure 1). After the 40 µm movement in x-direction, the stage moved 1 µm in the y-direction at maximum speed (speed factor 1). The movement in y-direction should also be optimized depending on the crystal size and distribution on the grid. The stage then returned -40 µm in x-direction at the original speed, followed by another 1 µm movement in y-direction. The final line repeats the initial four lines, and the stage sweeping continues until manually stopped in the *SerialEM* interface. It is important to note that similar procedures can be performed using any software that offers stage control and are not limited to *SerialEM*.

**c-SerialED Data Processing**

Processing, including data conversion, peak finding, beam center refinement, indexing, integration, and merging, is carried out using a customized workflow that combines in-house Python scripts with existing algorithms, such as peakfinder8 from *Cheetah*[6] and indexing and integration tools from the *CrystFEL* suite[7]. Each dataset was processed using a separate Jupyter-notebook file, which contains the processing parameters specific to the dataset presented in this study. In the case of lysozyme, there was a number of frames that were indexed with a unit cell parameter α far from the expected value. Therefore, a filter was applied to exclude crystals with a unit cell parameter α greater than 90°. Data reduction statistics for all c-SerialED datasets are summarized in Extended Data Table 3.

Data processing is done using custom python code as well as readily available software. Each dataset is processed in its individual Jupyter-Notebook, which contains a cell with function definitions and cells for parameter definitions for each individual processing step (Extended Data Figure 5).

In an initial step, output data from different data collection software is converted to HDF5 format. The code is capable of handling different input formats (*.emd* and *.ser*). Original frame dimensions and bit depth are kept; however, the newly written HDF5 image stack is chunked to improve I/O performance. Peak finding is done using *peakfinder8* implemented in Robert Bücker's *diffractem* package.[8] Extensive documentation for the use of this package can be found online. Peak information is stored in CXI format in the *.h5* files.[9] A two-step process of center estimation and refinement is used to determine the exact position of the beam behind the beam stop. By analyzing diffraction patterns for Friedel pairs and applying linear regression, the algorithm estimates the beam center for batches of subsequent frames in the data stack. These are interpolated to estimate the position for each individual frame. Subsequently the positions are iteratively refined by smoothing out deviations over the whole dataset using a *Locally Weighted Scatterplot Smoothing* (LOWESS) algorithm.[10] Indexing is done using Gevorkov's *extended gradient descent algorithm for lattice finding* (XGANDALF), part of CrystFEL.[11,12] Due to XGANDALF's high sensitivity to precise beam-center positioning, indexing is performed multiple times with systematically varied x and y beam-center offsets. A Python wrapper aggregates results across these runs, selecting the best-indexed frames, significantly improving the overall indexing rate. The best results are determined and written to a *.sol* file, to be used as an input for further processing steps. If a substantial portion of frames exhibit multiple indexing solutions with similarly high numbers of indexed peaks and comparable indexing rates, ambiguity resolution is performed using CrystFEL's *ambigator* module to determine the correct orientation.[13] The integrated *.stream* files are merged using CrystFEL's *partialator* module and converted to *.mtz* format using *get_hkl*.

The code presented here demonstrates a functional implementation of the workflow. An optimized, comprehensive processing suite packaged as a Python library, complemented by a graphical user interface, is currently under development and will be published separately.

### MicroED Data Collection and Processing

The EPUD software (Thermo Fisher Scientific) was utilized to collect atlas maps and MicroED data. MicroED data for DtpAa and MTH1 were collected with specifics as listed in Extended Data Table 2.

Data processing of all MicroED datasets was conducted using XDS, and the datasets were scaled and merged using XSCALE.[14] Data reduction statistics for the MicroED dataset can be found in Extended Data Table 4.

### Refinement and Structural Analysis

All structures were solved using molecular replacement in *Phaser*[15], using the following search models: PDB ID: 7SKW (lysozyme), 6TB8 (DtpAa-apo and DtpAa-N3), 4D12 (UOX–urate and UOX–9-methylurate), and 3ZR1 (MTH1–8DG). For the MicroED datasets, the DtpAa-apo structure was solved using the model derived from the c-SerialED DtpAa-apo structure, while both the high- and low-fluence MTH1–8DG datasets were solved using PDB ID: 3ZR1. Structure refinement was performed using *phenix.refine*[16]. Refinement statistics are listed in Extended Data Table 3 (c-SerialED) and Extended Data Table 4 (MicroED). For DtpAa, estimated bond length errors were calculated from the coordinate diffraction precision index (DPI) using the online server *online_DPI*[17].

In both the DtpAa and MTH1 structures, the A chain was better resolved than the B chain, likely due to differences in crystal packing. Consequently, analysis of the heme group in DtpAa and the bound 8-oxo-dGTP in MTH1 was restricted to the A chain. This observation is consistent with previous studies on these crystal forms.[18,19]

### Identification of Candidate Hydrogen atoms

Hydrogen analysis was performed using a method similar to that described by Clabbers et al.[20], with the modification that only peaks in the Fourier difference map (Fo–Fc) appearing above a threshold of 3 rmsd were considered as candidate hydrogen atoms to ensure they were above the noise level. Positive Fourier difference peaks were identified at positions that were likely to be a hydrogen atom. These peaks represent location of possible "candidate hydrogen atoms". Candidate hydrogen atoms were identified using the Fo-Fc difference map, calculated by *Phenix.refine*. A structure were all hydrogens had been deleted was used to generate a calculated map. Subsequently, *PEAKMAX* in the CCP4 software package was utilized to list peaks above a threshold of 3 rmsd. These peaks were manually inspected in Coot, and only those located in chemically reasonable positions were assigned as candidate hydrogen atoms. As a result, 41.93% (226) peaks were assigned as candidate hydrogen atoms. These atoms account for 21.6% of all non-water hydrogen atoms in the entire structure. After refinement, 225 candidate hydrogen atoms exhibited non-zero occupancy. Several representative types of candidate hydrogen atoms are shown in Extended Data Figure 3. The distribution of bond lengths for different types of hydrogens is presented in Extended Data Figure 4. Detailed information on all identified candidate H atoms is provided in the Supplementary Information (Extended Data Table 5-12).

Although several candidate hydrogens were identified, many non-water hydrogens in the structure could not be directly observed. To investigate whether information about hydrogens was still present, even when they could not be directly identified in the map, we performed an alternative analysis. In this approach, theoretical hydrogens were added for all atoms in the structure, irrespective of whether they were observed in the map, and the occupancies of these were refined alongside the other refinement parameters. In this global refinement of hydrogen occupancy, 143 out of the total 1028 non-water hydrogen atoms were refined to an occupancy of zero.

### Data Availability

The c-SerialED datasets have been deposited in the SBGrid Data Bank and are accessible via the following DOIs: Lysozyme (https://doi.org/10.15785/SBGRID/1150), DtpAa-apo (https://doi.org/10.15785/SBGRID/1149), DtpAa-N3 (https://doi.org/10.15785/SBGRID/1146), UOX–urate (https://doi.org/10.15785/SBGRID/1157), UOX–9-methylurate (https://doi.org/10.15785/SBGRID/1154), and MTH1–8DG (https://doi.org/10.15785/SBGRID/1163).

MicroED datasets are available on Zenodo for DtpAa-apo (https://doi.org/10.5281/zenodo.13640518), and MTH1–8DG (high fluence: https://doi.org/10.5281/zenodo.15168593; low fluence: https://doi.org/10.5281/zenodo.15168593).

### Code Availability

The Jupyter-notebook files used for data processing of the c-SerialED datasets are available on Zenodo (http://doi.org/10.5281/zenodo.15185823).

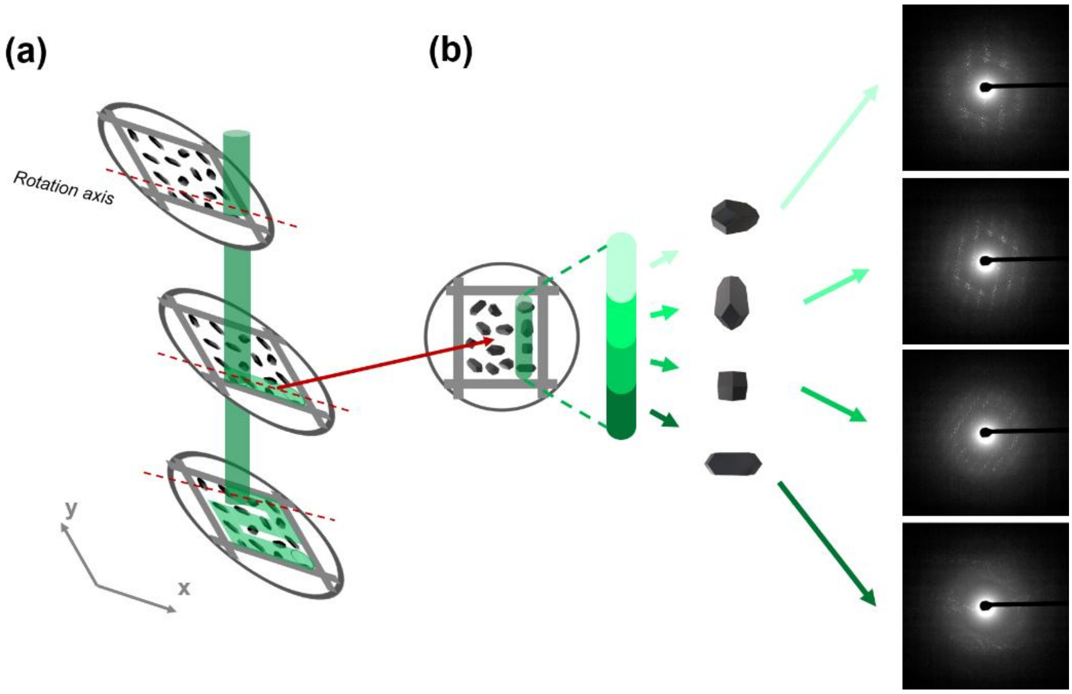


**Extended Data Figure 1.** Schematic illustration of data collection for c-SerialED data. a) Data were collected by stage moving. b) Diffraction patterns with different orientations were obtained during data collection.

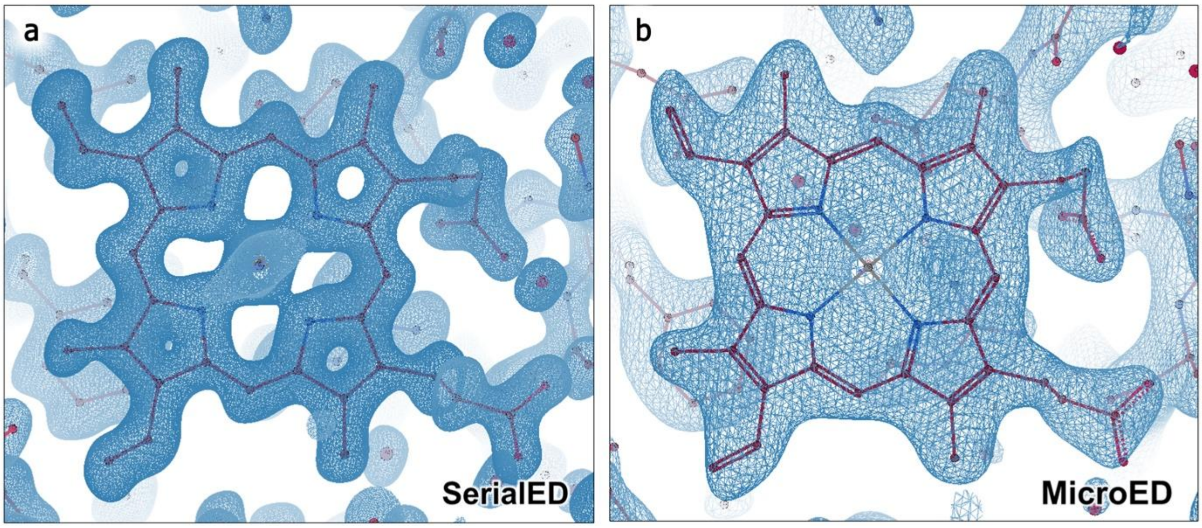


**Extended Data Figure 2.** Model and 2Fo-Fc maps of the chain A heme site in DtpAa-apo as resolved by (a) c-SerialED, refined to 1.3 Å, and (b) MicroED, refined to 2.5 Å. The maps are shown as non-filled 2Fo-Fc maps contoured at 1.5 rmsd (blue mesh).

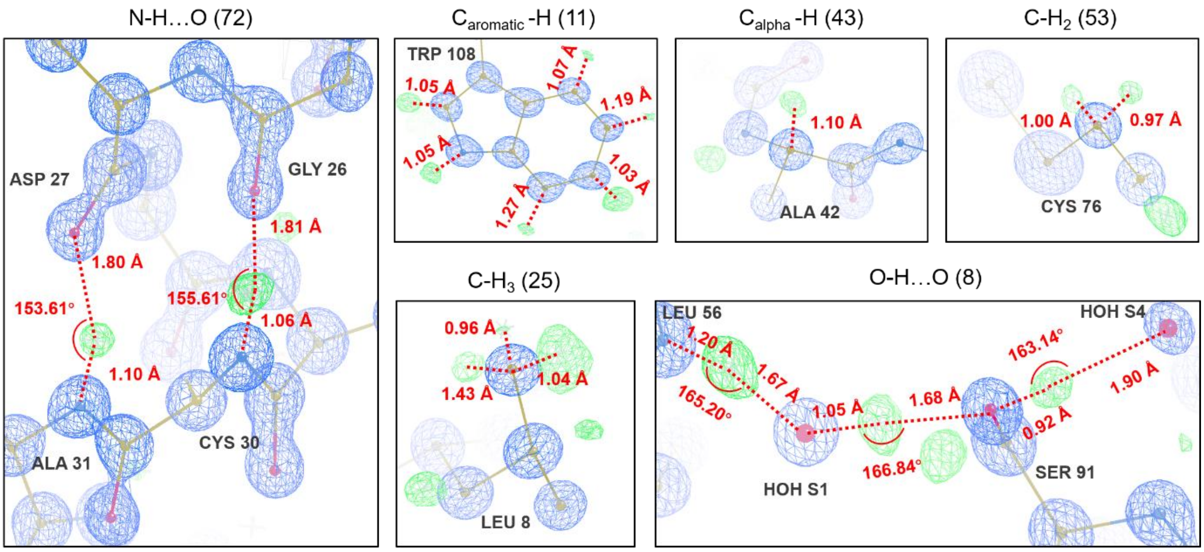


**Extended Data Figure 3.** Examples of selected hydrogen candidates in the lysozyme structure. The types of hydrogen atoms are labeled above each panel, with the number of observed peaks for each hydrogen type indicated in parentheses. Bond lengths and bond angles are marked in each image. The unfilled 2Fo–Fc electrostatic potential map (blue) is contoured at 4 rmsd, while the Fo–Fc difference map (green and red) is contoured at 2.9–3.5 rmsd for improved inspection.

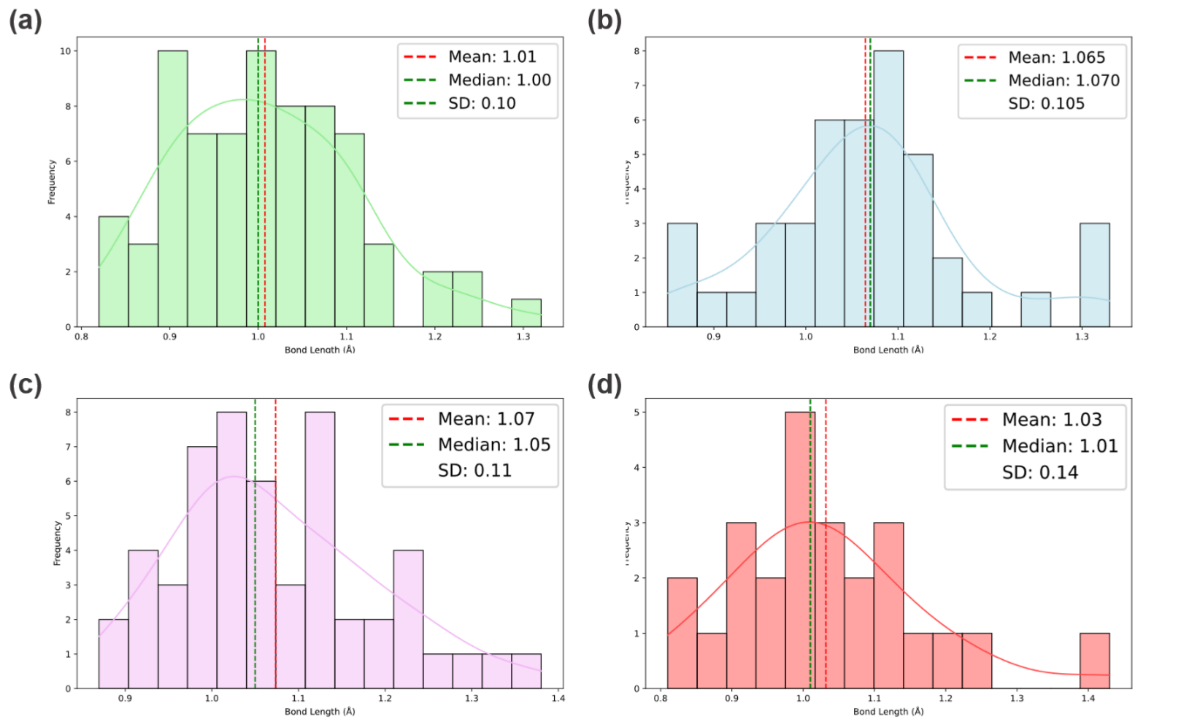


**Extended Data Figure 4.** Distribution of bond lengths for candidate hydrogen atom types with more than 20 observations in the lysozyme structure: (a) N–H···O, (b) Cα–H, (c) $C–H_2$, and (d) $C–H_3$. Mean bond lengths, medians, and standard deviations are indicated in the upper right corner of each panel.

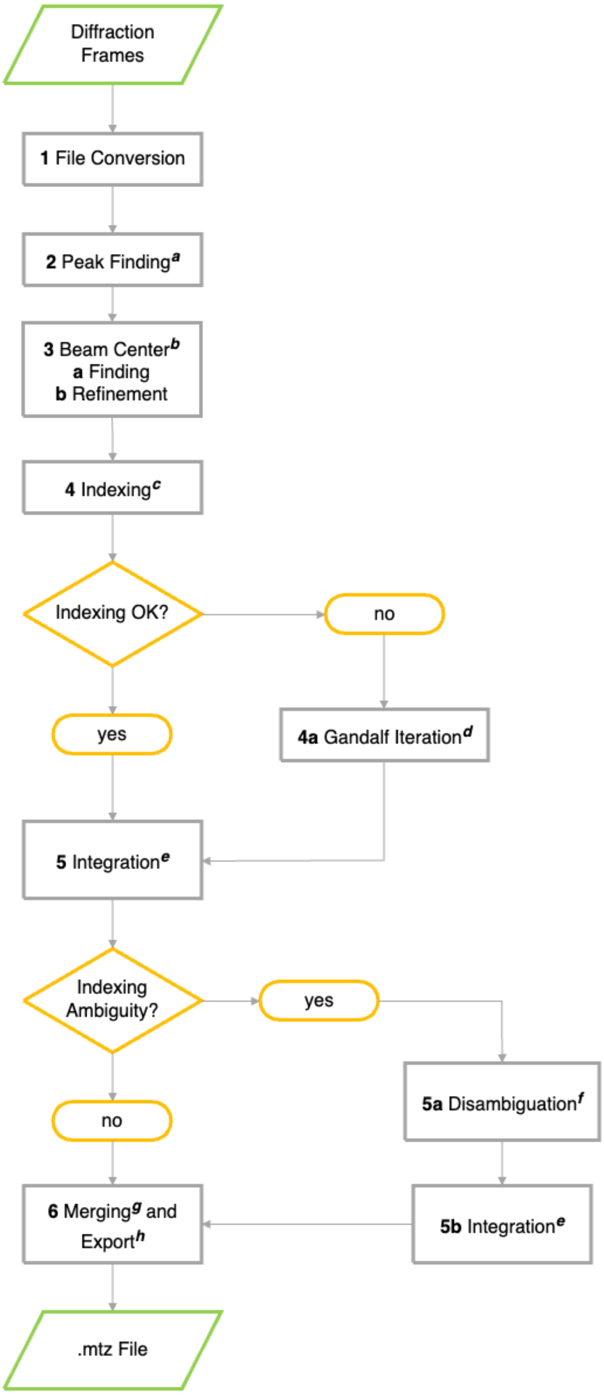


**Extended Data Figure 5.** Data processing flowchart (a utilizes peakfinder8 as part of diffractem, b custom algroithm to determine beam centers based on Friedel pairs, c indexing using xgandalf implemented in CrystFEL's indexamajig, d indexing using a custom wrapper for xgandalf/indexamajig to iterate over subpixel beam center offsets, e integration using xgandalf implemented in CrystFEL's indexamajig, f resolution of indexing ambiguities using CrystFEL's ambigator module, g merging of intensities using CrystFEL's partialator module, h export to .mtz format (CCP4) using CrystFEL's get_hkl module)

**Extended Data Table 1.** c-SerialED data collection details.

| | **Camera length [m]** | **C2 aperture [μm]** | **Spot size [-]** | **Beam size [nm]** | **Stage speed [μm $s^{-1}$]** | **Stage tilt [°]** | **Exposure time [ms]** | **Fluence per frame** [$e^{-}Å^{-2}$] | **No. frames integrated [-]** | **Resolution of final dataset [Å]** |
|---|---|---|---|---|---|---|---|---|---|---|
| Lysozyme | 0.68 | 20 | 4 | 300 | 3.1 | 45 | 52.6 | 1.52 | 30 807 | 11.73–0.83 |
| DtpAa-apo | 1.1 | 20 | 4 | 500 | 4.7 | 0 | 52.6 | 2.2 | 21 036 | 19.03–1.30 |
| DtpAa-N3 | 1.1 | 20 | 4 | 300 | 3.1 | 0 | 52.6 | 3.5 | 46 423 | 21.47–1.10 |
| UOX-urate | 1.35 | 20 | 4 | 450 | 4.2 | 0 | 72 | 2.2 | 3 823 | 26.38–1.75 |
| UOX-9-methylurate | 1.1 | 20 | 5 | 450 | 4.2 | 0 | 72 | 1.2 | 56 538 | 19.82–1.25 |
| MTH1 | 1.35 | 20 | 2 | 300 | 8.3 | 45 | 52.6 | 5.63 | 10 801 | 25.73–1.66 |

**Extended Data Table 2.** MicroED data collection details.

| | **Camera length [m]** | **C2 aperture [µm]** | **Spot size [-]** | **Beam size [µm]** | **Tilt per frame [°]** | **Tilt range [°]** | **Exposure time [s]** | **Fluence [e⁻Å⁻²]** | **No. merged datasets [-]** | **Resolution final dataset [Å]** |
|---|---|---|---|---|---|---|---|---|---|---|
| DtpAa-apo | 1.2 | 20 | 9 and 8* | 1.5 and 3* | 1 | 40-70 | 1 | 0-5.0 | 20 | 20–2.5 |
| MTH1-high-dose | 1.7 | 20 | 7 | 1.5 | 1 | 20 | 1 | 0-7.4 | 20 | 29.35–2.32 |
| MTH1-low-dose | 1.7 | 20 | 10 | 1.5 | 1 | 20 | 1 | 0-1.1 | 25 | 29.67–2.86 |

* Two datasets of DtpAa were collected using a wider beam of 3 µm beam and spot size 8, with fluence measured to 0.210 eÅ-2. The broader beam was employed to mitigate the risk of crystal loss at high tilt angles.

**Extended Data Table 3.** Data reduction and refinement statistics for structures refined against c-SerialED data.

| **Dataset** | **Lysozyme** | **DtpAa-apo** | **DtpAa-N3** | **UOX-9-methyl-urate** | **UOX-urate** | **MTH1-8DG** |
|---|---|---|---|---|---|---|
| Cumulative Fluence [eÅ$^{-2}$] | 0-1.52 | 0-3.5 | 0-3.5 | 0-1.2 | 0-2.2 | 0-5.63 |
| Number of integrated frames* | 34,231 | 21,036 | 41,986 | 56,538 | 3,823 | 10,801 |
| Space Group | *P*1 | *P*2$_1$ | | *I*222 | *I*222 | *P*2$_1$2$_1$2$_1$ |
| Unit cell | | | | | | |
| *a; b; c* [Å] *α; β; γ* [°] | 26.66; 31.15; 33.57; 87.73; 108.97; 111.6 | 72.77; 67.4; 73.63; 90; 105.8; 90 | | 80.58; 94.49; 103.89; 90; 90; 90 | 80.13; 94.44; 103.97; 90; 90; 90 | 59.34; 67.55; 80.11; 90; 90; 90 |
| Resolution [Å]** | 11.73–0.83 (0.86–0.83) | 19.03–1.30 (1.35–1.30) | 21.47–1.10 (1.14–1.10) | 19.82–1.25 (1.29–1.25) | 26.38–1.75 (1.81–1.75) | 25.73–1.66 (1.72–1.66) |
| No. reflections (total/unique) | 5,003,763/ 89,374 | 8,818,292/ 137,391 | 15,635,343/ 250,678 | 24,688,447/ 109,263 | 801,293/ 38,877 | 3,882,602/ 38,744 |
| Multiplicity** | 55.99 (57.2) | 64.2 (69.0) | 62.4 (68.1) | 225.95 (106.99) | 20.61 (14.6) | 100.21 (34.3) |
| *I/SigI*** | 4.55 (0.46) | 4.72 (0.5) | 3.73 (0.26) | 5.76 (0.88) | 4.28 (1.54) | 5.27 (0.71) |
| $R_{split}$** [%] | 0.109 (2.594) | 0.164 (1.867) | 0.195 (5.105) | 0.127 (1.359) | 0.219 (0.936) | 0.136 (1.873) |
| $CC_{1/2}$** | 0.994 (0.28) | 0.988 (0.406) | 0.988 (0.301) | 0.995 (0.393) | 0.962 (0.392) | 0.986 (0.210) |
| $CC_{star}$** | 0.998 (66.2) | 0.997 (0.760) | 0.997 (0.680) | 0.999 (0.751) | 0.99 (0.75) | 0.996 (0.589) |
| Completeness [%]** | 98.3 (87.5) | 80.02 (72.34) | 79.62 (33.06) | 99.96 (99.11) | 96.93 (95.96) | 96.4 (65.4) |
| No. reflections used in refinement** | 87,855 (7,846) | 134,402 (12,091) | 220,288 (9,103) | 109,106 (10,727) | 32,397 (5,244) | 37,120 (2,467) |
| $R_{work}/R_{free}$ | 0.1755/0.2064 | 0.2123/ 0.2473 | 0.2162/ 0.2391 | 0.1913/0.2031 | 0.2205/0.2429 | 0.2051/0.2528 |
| RMSD bond lengths [Å] | 0.019 | 0.0130 | 0.0104 | 0.011 | 0.006 | 0.010 |
| RMSD bond angles [°] | 1.22 | 1.07 | 0.99 | 1.1 | 0.642 | 0.947 |
| Ramachandran plot | | | | | | |
| Most favoured [%] | 99.21 | 98.48 | 98.48 | 98.27 | 97.6 | 99.03 |
| Allowed [%] | 0.79 | 1.52 | 1.52 | 1.73 | 2.4 | 0.97 |
| Outliers [%] | 0 | 0 | 0 | 0 | 0 | 0 |
| Rotamer outliers [%] | 0.91 | 0.54 | 0.36 | 0.67 | 0.34 | 0 |
| PDB accession code | 9QUM | 9FYK | 9FY7 | 9QW5 | 9QW6 | 9QUE |

* Number of merged patterns in CrystFEL.
** Values in parentheses are for the highest resolution shell.

**Extended Data Table 4.** Data reduction and refinement statistics for structures refined against MicroED data.

| Dataset | DptAa-apo | MTH1-8DG low-fluence | MTH1-8DG high-fluence |
|---|---|---|---|
| Cumulative Fluence [eÅ$^{-2}$] | 0-5 | 0-1.1 | 0-7.4 |
| Number of merged datasets | 20 | 25 | 20 |
| Space Group | $P2_1$ | $P2_12_12_1$ | $P2_12_12_1$ |
| Unit cell | | | |
| *a; b; c* [Å] | 72.77; 67.4; 73.63 | 59.34; 67.55; | 59.34; 67.55; |
| *α; β; γ* [°] | 90; 105.8; 90 | 80.11; 90; 90; 90 | 80.11; 90; 90; 90 |
| Resolution [Å]** | 19.46–2.40 (2.46 –2.40) | 29.63–2.86 (3.02–2.86) | 29.35–2.32 (2.40–2.32) |
| Number of reflections (total/unique) | 456,381/48,362 | 127,640/6,638 | 152,619/11,420 |
| Multiplicity* | 10.6 (10.0) | 19.2 (10.4) | 13.4 (8.4) |
| *I/SigI** | 3.42 (1.25) | 5.8 (1.7) | 5.5 (1.6) |
| $R_{meas}$* [%] | 55.3 (183.0) | 55.9 (159.5) | 47.6 (163.8) |
| $CC_{1/2}$* | 90.3 (27.8) | 91.3 (43.6) | 97.4 (48.6) |
| $CC_{star}$* | - | - | - |
| Completeness [%]* | 87.8 (79.3) | 84.1 (53.9) | 77.7 (45.0) |
| Number of reflections used in refinement* | 23,683 (2,099) | 6,601 (411) | 11,355 (635) |
| $R_{work}/R_{free}$ | 0.2020/0.2564 | 0.2314/0.2824 | 0.2036/0.2624 |
| RMSD bond lengths [Å] | 0.0026 | 0.002 | 0.002 |
| RMSD bond angles [°] | 0.56 | 0.478 | 0.489 |
| Ramachandran plot | | | |
| Most favoured [%] | 97.10 | 94.43 | 96.07 |
| Allowed [%] | 2.90 | 5.57 | 3.93 |
| Outliers [%] | 0 | 0 | 0 |
| Rotamer outliers | 0.36 | 1.47 | 1.47 |
| PDB accession code | 9FYH | 9QUK | 9QUH |

* Values in parentheses are for the highest resolution shell.

**Extended Data Table 5.** Peak height of the difference peak in the Fc–Fo map for a candidate hydrogen on a nitrogen donor. The peak height is reported as the contour level (rmsd) at which the peak is no longer visible. The table also includes the corresponding hydrogen donor, the donor–hydrogen (D–H) bond distance, the donor–acceptor (D···A) distance, and the D–H···A angle. The atom label and residue of the acceptor are also provided. The letters "A" and "B" in parentheses indicate split occupancy of amino acid residues. Total observations: 72.

| | | | From Q peaks | | | After refinement | | |
|---|---|---|---|---|---|---|---|---|
| **Peak rmsd** | **Donator (Atom-Residue)** | **Acceptor (Atom-Residue)** | **D-H (Å)** | **D···A (Å)** | **D-H···A (◦ )** | **D-H (Å)** | **D···A (Å)** | **D-H···A (◦ )** |
| 3.93 | N-4 GLY | O-S132 | 1.32 | 1.67 | 162.81 | 0.87 | 2.12 | 160.72 |
| 3.34 | N-5 ARG | O-S92 | 1.13 | 1.99 (A) | 176.72 (A) | 0.86 | 2.25 | 157.51 |
| 3.63 | NH1-5 ARG | O-123 TRP | 1.11 | 1.89 (A) | 135.05 (A) | 0.86 | 2.27 | 145.96 |
| 3.25 | N-7 GLU | OE1-7 | 0.85 | 1.9 | 148.09 | 0.86 | 1.91 | 150.10 |
| 3.07 | N-8 LEU | O-4 GLY | 1.06 | 1.87 | 141.13 | 0.87 | 1.97 | 157.63 |
| 3.92 | N-9 ALA | O-5 ARG | 1.00 | 1.86 (A) | 170.02 (A) | 0.88 | 2.02 | 164.40 |
| 3.95 | N-11 ALA | O-7 GLU | 1.11 | 1.94 | 151.68 | 0.87 | 2.13 | 161.22 |
| 3.25 | N-12 MET | O-8 LEU | 0.91 | 1.78 | 176.44 | 0.86 | 1.86 | 174.04 |
| 4.15 | N-13 LYS | O-9 ALA | 0.99 | 1.76 | 167.76 | 0.87 | 1.91 | 161.75 |
| 3.54 | N-14 ARG | O-10 ALA | 1.01 | 1.9 | 153.28 | 0.86 | 2.04 | 158.48 |
| 3.6 | N-15 HIS | O-11 ALA | 0.91 | 2.23 | 140.55 | 0.86 | 2.27 | 141.92 |
| 4.61 | N-17 LEU | O-12 MET | 1.09 | 1.99 | 161.94 | 0.87 | 2.20 | 174.02 |
| 3.69 | N-18 ASP | O-S3 HOH | 0.96 | 1.94 | 174.74 | 0.86 | 2.06 | 163.44 |
| 3.39 | N-19 ASN | O-23 TYR | 1 | 2.06 | 131.51 | 0.87 | 2.11 | 137.98 |
| 3.25 | ND2-19 ASN | O-81 SER | 1.08 | 1.8 | 177.78 | 0.87 | 2.02 | 176.02 |
| 4.08 | N-20 TYR | O-17 LEU | 1.03 | 1.95 | 158.52 | 0.87 | 2.10 | 174.34 |
| 3.25 | N-21 ARG | O-S8 HOH | 0.97 | 2.02 | 129.31 | 0.86 | 1.96 | 154.83 |
| 3.08 | N-22 GLY | O-19 ASN | 0.96 | 2.08 | 174.21 | 0.87 | 2.19 | 169.76 |
| 4.26 | N-24 SER | O-S12 | 1.11 | 1.78 | 171.59 | 0.87 | 2.05 | 163.57 |
| 3.69 | N-25 LEU | OD1-18 | 0.96 | 1.89 | 172.95 | 0.87 | 2.13 | 141.90 |
| 4 | N-29 VAL | O-25 LEU | 0.88 | 2.25 | 159.6 | 0.86 | 2.29 | 161.49 |
| 4.56 | N-30 CYS | O-26 GLY | 1.06 | 1.81 | 155.61 | 0.88 | 1.96 | 168.59 |
| 4.29 | N-31 ALA | O-27 ASN | 1.1 | 1.8 | 153.61 | 0.87 | 2.02 | 155.75 |
| 3.74 | N-32 ALA | O-28 TRP | 0.89 | 2.12 | 155.63 | 0.86 | 2.13 | 164.55 |
| 3.07 | N-35 GLU | O-31 ALA | 0.91 | 1.97 | 142.1 | 0.87 | 1.99 | 145.51 |
| 3.3 | N-37 ASN | O-33 LYS | 1.08 | 2.01 | 127.86 | 0.86 | 2.10 | 139.34 |
| 3.19 | N-38 PHE | O-32 ALA | 1.08 | 2.33 | 166.27 | 0.87 | 2.55 | 162.61 |
| 3.19 | N-40 THR | O-1 LYS | 1.23 | 1.73 | 140 | 0.87 | 1.94 | 164.71 |
| 3.35 | N-41 GLN | OD1-39 | 1.08 | 1.82 (A) | 159.93 (A) | 0.88 | 1.97 | 162.58 |
| 3.87 | N-42 ALA | O-39 ASN | 0.94 | 2.16 | 172.17 | 0.86 | 2.23 | 171.13 |
| 4.34 | N-44 ASN | O-52 ASP | 0.94 | 2 | 168.7 | 0.87 | 2.10 | 153.25 |
| 3.78 | NH2-45 ARG | O-S70 | 1.04 | 1.87 | 149.31 | 0.86 | 2.01 | 136.73 |
| 3.44 | NH2-45 ARG | O-S63 | 1.2 | 1.79 | 166 | 0.86 | 2.16 | 155.36 |
| 3.16 | N-49 GLY | O-46 ASN | 0.97 | 2.46 | 144.75 | 0.86 | 2.48 | 162.13 |
| 4.11 | N-50 SER | OD1-48 | 0.86 | 2.11 | 150.49 | 0.86 | 2.05 | 165.77 |
| 3.54 | N-51 THR | OG-60 | 0.94 | 1.93 | 165.81 | 0.86 | 2.02 | 162.51 |
| 4.01 | N-52 ASP | O-44 ASN | 1.03 | 1.77 | 173.95 | 0.87 | 1.96 | 163.85 |

| | | | | | | | | |
|---|---|---|---|---|---|---|---|---|
| 4.53 | N-55 ILE | O-S2 HOH | 1.01 | 1.92 | 147.4 | 0.87 | 2.07 | 144.49 |
| 4.46 | N-56 LEU | O-S1 HOH | 1.2 | 1.67 | 165.2 | 0.87 | 2.00 | 164.27 |
| 3.52 | N-57 GLN | O-54- | 0.95 | 1.99 | 152.36 | 0.86 | 2.08 | 154.86 |
| 3.63 | NE2-57 GLN | O-54 GLY | 1.13 | 1.71 | 167.5 | 0.87 | 1.99 | 171.01 |
| 3.87 | N-58 ILE | O-53 TYR | 0.9 | 2.08 | 161.46 | 0.87 | 2.14 | 157.93 |
| 3.09 | N-59 ASN | O-201 | 0.91 | 1.97 | 152.29 | 0.86 | 2.02 | 156.57 |
| 3.53 | N-60 SER | O-51 THR | 1.08 | 1.9 | 153.37 | 0.86 | 2.14 | 147.03 |
| 3.52 | N-61 ARG | OD1-59 | 0.82 | 2.08 | 161.38 | 0.85 | 2.05 | 169.13 |
| 3.06 | N-62 TRP | O-S153 | 0.85 | 2.05 | 149.96 | 0.86 | 2.13 | 136.87 |
| 3.95 | N-63 TRP | O-59 ASN | 0.9 | 2.21 | 156.1 | 0.85 | 2.22 | 163.09 |
| 4.25 | N-65 ASN | O-78 ILE | 0.91 | 1.97 | 169.52 | 0.86 | 2.04 | 164.37 |
| 3.26 | N-66 ASP | O-S11 | 1.1 | 1.77 | 159.01 | 0.86 | 2.02 | 160.38 |
| 3.08 | N-68 ARG | OD1-66 | 1.15 | 1.92 | 136.31 | 0.86 | 2.04 | 164.68 |
| 3.01 | N-73 ARG | O-61 ARG | 1 | 2.06 | 134.87 | 0.86 | 2.05 | 154.07 |
| 3.29 | N-75 LEU | O-62 TRP | 0.83 | 1.98 | 165.73 | 0.85 | 1.96 | 167.54 |
| 4.26 | N-76 CYS | O-63 TRP | 0.95 | 1.98 | 139.73 | 0.87 | 1.96 | 156.32 |
| 3.06 | ND2-77 ASN | O-S32 | 1.11 | 2.02 | 151.36 | 0.86 | 2.28 | 177.05 |
| 3.02 | N-78 ILE | OD1-74 | 0.98 | 1.99 (A) | 154.58 (A) | 0.86 | 2.11 | 156.59 |
| 3.68 | N-82 ALA | O-79 PRO | 1.08 | 1.89 | 159.81 | 0.86 | 2.10 | 158.82 |
| 5.8 | N-83 LEU | O-80 CYS | 0.96 | 1.96 | 152.36 | 0.87 | 2.04 | 154.46 |
| 5 | N-84 LEU | O-81 SER | 0.93 | 2.08 | 169.51 | 0.87 | 2.14 | 168.44 |
| 3.1 | N-91 SER | O-S4 HOH | 1.05 | 2.18 | 129.25 | 0.86 | 2.25 | 138.96 |
| 3.97 | N-92 VAL | O-88 ILE | 1 | 1.91 | 145.82 | 0.87 | 1.99 | 157.99 |
| 3.35 | N-93 ASN | O-89 THR | 0.9 | 1.89 | 169.01 | 0.86 | 1.93 | 171.31 |
| 4.3 | N-94 CYS | O-90 ALA | 0.86 | 2.02 | 167.24 | 0.86 | 2.03 | 164.27 |
| 3.26 | N-97 LYS | O-93 ASN | 1.02 | 2.1 | 151.78 | 0.87 | 2.24 | 155.05 |
| 4.15 | N-99 VAL | O-95 ALA | 0.92 | 2.21 | 156.11 | 0.87 | 2.26 | 158.51 |
| 3.46 | NE1-108 TRP | O-56 LEU | 1.05 | 1.77 | 154.62 | 0.86 | 2.11 | 131.54 |
| 3.69 | N-109 VAL | O-S14 | 1.03 | 2.04 | 135.06 | 0.86 | 2.07 | 158.76 |
| 3.36 | N-113 ASN | O-109 | 1 | 1.91 | 162.71 | 0.87 | 2.07 | 157.66 |
| 4.05 | NH2-114 ARG | O-S41 | 1.02 | 1.97 | 162.9 | 0.86 | 2.28 | 147.37 |
| 3.7 | N-116 LYS | O-118 TRP | 1.01 | 1.72 | 174.3 | 0.87 | 1.88 | 169.31 |
| 3.66 | N-118 THR | O-115 | 1.25 | 1.79 (A) | 151.86 (A) | 0.86 | 2.21 | 144.19 |
| 3.73 | N-123 TRP | O-120 | 0.91 | 2.15 | 138 | 0.86 | 2.03 | 166.30 |
| 3.22 | N-126 GLY | O-S50 | 0.99 | 1.73 | 153.82 | 0.86 | 1.81 | 169.75 |

**Extended Data Table 6.** Peak height of the difference peak in the Fc–Fo map for a candidate hydrogen on an oxygen donor. The peak height is reported as the contour level (rmsd) at which the peak is no longer visible. The table also includes the corresponding hydrogen donor, the donor–hydrogen (D–H) bond distance, the donor–acceptor (D···A) distance, and the D–H···A angle. The atom label and residue of the acceptor are also provided. Total observations: 8.

| | | | From Q peaks | | | After refinement | | |
|---|---|---|---|---|---|---|---|---|
| **Peak rmsd** | **Donator (Atom-Residue)** | **Acceptor (Atom-Residue)** | **D-H (Å)** | **D···A (Å)** | **D-H···A (◦ )** | **D-H (Å)** | **D···A (Å)** | **D-H···A (◦ )** |
| 3.53 | OG-36 SER | O-55 ILE | 0.83 | 1.96 | 152.31 | 0.84 | 1.91 | 162.84 |
| 3.91 | OH-53 TYR | OD2-66 | 1.04 | 1.61 | 162.81 | 0.85 | 1.79 | 167.94 |
| 3.29 | OG-60 SER | OG1-69 | 0.95 | 1.74 | 148.39 | 0.84 | 1.80 | 160.65 |
| 3.89 | OG-91 SER | O-S4 HOH | 0.92 | 1.9 | 163.14 | 0.84 | 1.90 | 169.46 |
| 4.11 | O-S1 HOH | OG-91 SER | 1.05 | 1.68 | 166.84 | H of waters not modeled | | |
| 3.64 | O-S132 | O-S134 | 1.18 | 1.73 | 137.46 | | | |
| 3.17 | O-S3 HOH | O-13 LYS | 1.28 | 1.54 | 154.65 | | | |
| 3.08 | O-S8 HOH | O-S39 | 0.88 | 1.77 | 161.93 | | | |

**Extended Data Table 7.** Peak height of the difference peak in the Fc–Fo map for a candidate hydrogen on an aromatic carbon ($C_{aromatic}$). The peak height is reported as the contour level (rmsd) at which the peak is no longer visible. The table also includes the atom label and residue of the binding atom, as well as the bond distance between the binding atom and the candidate hydrogen (Atom–H). Total observations: 11.

| Peak rmsd | Binding atom | From Q peaks Atom-H (Å) | After refinement Atom-H (Å) |
|---|---|---|---|
| 3.64 | CD1-23 TYR | 0.99 | 0.93 |
| 3.51 | CZ3-28 TYR | 0.87 | 0.93 |
| 3.34 | CE3-28 TYR | 0.99 | 0.93 |
| 3.06 | CZ2-28 TYR | 1.32 | 0.93 |
| 3.23 | CZ-38 PHE | 0.96 | 0.93 |
| 3.8 | CE1-53 TYR | 0.98 | 0.94 |
| 4.18 | CH2-108 TRP | 1.03 | 0.93 |
| 4.03 | CD1-108 TRP | 1.05 | 0.94 |
| 3.09 | CZ2-108 TRP | 1.27 | 0.93 |
| 3.06 | CE3-108 TRP | 1.07 | 0.93 |
| 3.04 | CZ3-108TRP | 1.19 | 0.93 |

**Extended Data Table 8.** Peak height of the difference peak in the Fc–Fo map for a candidate hydrogen on an α-carbon ($C_\alpha$). The peak height is reported as the contour level (rmsd) at which the peak is no longer visible. The table also includes the atom label and residue of the binding atom, as well as the bond distance between the binding atom and the candidate hydrogen (Atom–H). Total observations: 43.

| Peak rmsd | Binding atom | From Q peaks Atom-H (Å) | After refinement Atom-H (Å) |
|---|---|---|---|
| 3.51 | CA-5 ARG | 1.05 (A) | 1.02 (A) |
| 3.33 | CA-7 GLU | 1.01 | 0.98 |
| 4.22 | CA-10 ALA | 1.16 | 0.98 |
| 3.13 | CA-13 LYS | 1.11 | 0.97 |
| 4.17 | CA-14 ARG | 0.97 | 0.97 |
| 3.75 | CA-20 TYR | 1.1 | 0.98 |
| 4.19 | CA-23 TYR | 1.18 | 0.99 |
| 3.07 | CA-25 LEU | 1.1 | 0.98 |
| 3.23 | CA-28 TRP | 1 | 0.98 |
| 3.23 | CA-31 ALA | 1.07 | 0.98 |
| 4.17 | CA-32 ALA | 0.97 | 0.96 |
| 3.37 | CA-40 THR | 1.11 | 0.99 |
| 3.77 | CA-42 ALA | 1.1 | 0.96 |
| 3.38 | CA-45 ARG | 0.85 | 0.97 |
| 3.21 | CA-46 ASN | 1.33 | 0.97 |
| 3.12 | CA-51 THR | 1.03 | 0.98 |
| 3.82 | CA-52 ASP | 1.03 | 0.98 |
| 3.02 | CA-54 GLY | 0.86 | 0.97 |
| 4.27 | CA-56 LEU | 0.94 | 0.97 |
| 3.55 | CA-57 GLN | 1.03 | 0.99 |
| 3.82 | CA-58 ILE | 0.91 | 0.98 |
| 4.07 | CA-59 ASN | 1.03 | 0.98 |
| 4.2 | CA-60 SER | 0.97 | 0.96 |
| 3.4 | CA-63 TRP | 1.08 | 0.98 |
| 3.41 | CA-74 ASN | 1.3 | 0.98 |
| 3.58 | CA-75 LEU | 1.09 | 0.97 |
| 4.38 | CA-76 CYS | 1.14 | 0.97 |
| 4.24 | CA-77 ASN | 1.06 | 0.98 |
| 5.26 | CA-80 CYS | 1.11 | 0.99 |
| 4.17 | CA-83 LEU | 0.87 | 0.97 |
| 3.79 | CA-88 ILE | 1.24 | 0.98 |
| 3.17 | CA-89 THR | 1.31 | 0.98 |
| 4.52 | CA-94 CYS | 1.12 | 0.98 |
| 3.37 | CA-95 ALA | 1 | 0.98 |
| 3.63 | CA-96 LYS | 1.08 | 0.98 |
| 3.17 | CA-98 ILE | 1.03 | 0.98 |
| 3.31 | CA-107 ALA | 1.08 | 0.98 |
| 3.43 | CA-111 TRP | 1.08 | 0.97 |
| 3.45 | CA-112 ARG | 1.11 | 0.97 |
| 3.12 | CA-113 ASN | 1.06 | 0.98 |

| | | | |
|---|---|---|---|
| 3.06 | CA-115 CYS | 1.06 | 0.98 |
| 4.42 | CA-116 LYS | 0.98 | 0.98 |
| 3.62 | CA-124 ILE | 1.07 | 0.97 |

**Extended Data Table 9.** Peak height of the difference peak in the Fc–Fo map for a candidate hydrogen on a tertiary carbon of the sidechain (C-H). The peak height is reported as the contour level (rmsd) at which the peak is no longer visible. The table also includes the atom label and residue of the binding atom, as well as the bond distance between the binding atom and the candidate hydrogen (Atom–H). Total observations: 10.

| **Peak rmsd** | **Binding atom** | **From Q peaks Atom-H (Å)** | **After refinement Atom-H (Å)** |
|---|---|---|---|
| 3.29 | CG-8 LEU | 1.1 | 0.97 |
| 3.13 | CB-29 VAL | 1.03 | 0.97 |
| 3.76 | CB-43 THR | 1.27 | 0.99 |
| 3.17 | CB-47 THR | 1.18 | 0.97 |
| 4.25 | CB-51 THR | 1.28 | 0.98 |
| 3.54 | CG-56 LEU | 1.07 | 0.98 |
| 3.51 | CB-58 ILE | 1.08 | 0.97 |
| 3.29 | CG-75 LEU | 1.12 | 0.98 |
| 3.64 | CB-88 ILE | 1.16 | 0.98 |
| 5.31 | CB-99 VAL | 1.32 | 0.97 |

**Extended Data Table 10.** Peak height of the difference peak in the Fc–Fo map for a candidate hydrogen on a nitrogen not involved in hydrogen bonding. The peak height is reported as the contour level (rmsd) at which the peak is no longer visible. The table also includes the atom label and residue of the binding atom, as well as the bond distance between the binding atom and the candidate hydrogen (Atom–H). Total observations: 4.

| **Peak rmsd** | **Binding atom** | **From Q peaks**<br>**Atom-H (Å)** | **After refinement**<br>**Atom-H (Å)** |
|---|---|---|---|
| 3.3 | N-67 GLY | 0.93 | 0.86 |
| 3.43 | N-71 GLY | 0.88 | 0.86 |
| 4.84 | N-88 ILE | 0.96 | 0.87 |
| 4.97 | N-117 GLY | 1.12 | 0.87 |

**Extended Data Table 11.** Peak height of the difference peak in the Fc–Fo map for a candidate hydrogen on a methylene carbon ($CH_2$). The peak height is reported as the contour level (rmsd) at which the peak is no longer visible. The table also includes the atom label and residue of the binding atom, as well as the bond distance between the binding atom and the candidate hydrogen (Atom–H). Total observations: 53.

| Peak rmsd | Binding atom | From Q peaks Atom-H (Å) | After refinement Atom-H (Å) |
|---|---|---|---|
| 3.95 | CG-1 LYS | 1.23 | 0.98 |
| 3.49 | CB-3 PHE | 0.98 | 0.97 |
| 3.3 | CB-7 GLU | 1.11 | 0.98 |
| 4.03 | CB-8 LEU | 0.91 | 0.97 |
| 3.46 | CG-12 MET | 1.14 | 0.98 |
| 3.23 | CG-14 ARG | 1.05 | 0.97 |
| 3.12 | CB-17 LEU | 1.01 | 0.98 |
| 3.12 | CB-18 ASP | 1.08 | 0.98 |
| 3.51 | CB-19 ASN | 1.03 | 0.97 |
| 3.45 | CA-22 GLY | 1.15 | 0.97 |
| 3.14 | CB-23 TYR | 1.2 | 0.98 |
| 3.04 | CB-25 LEU | 1.33 | 0.97 |
| 3.09 | CB-27 ASN | 1.01 | 0.98 |
| 3.55 | CB-30 CYS | 1.04 | 0.98 |
| 3.2 | CB-33 LYS | 1.23 | 0.97 |
| 3.26 | CB-35 GLU | 1.03 | 0.97 |
| 3.9 | CB-37 ASN | 1.07 | 0.97 |
| 5.49 | CB-41 GLN | 1.08 | 0.98 |
| 3.64 | CG-41 GLN | 0.99 | 0.97 |
| 4.42 | CB-45 ARG | 1.22 | 0.98 |
| 3.28 | CG-45 ARG | 0.99 | 0.97 |
| 3.16 | CB-46 ASN | 1.14 | 0.97 |
| 3.3 | CA-49 GLY | 1.03 | 0.97 |
| 3.52 | CB-50 SER | 1.27 | 0.97 |
| 3.75 | CG1-55 ILE | 1.14 | 0.97 |
| 3.28 | CB-56 LEU | 0.93 | 0.97 |
| 4.1 | CB-57 GLN | 0.89 | 0.97 |
| 3.78 | CG-57 GLN | 0.97 | 0.98 |
| 3.86 | CB-60 SER | 1.28 | 0.98 |
| 3.59 | CB-63 TRP | 1.04 | 0.97 |
| 3.45 | CB-64 CYS | 1.03 | 0.97 |
| 3.96 | CB-74 ASN | 1.03 | 0.97 |
| 3.1 | CB-74 ASN | 1.07 | 0.97 |
| 3.68 | CB-76 CYS | 1 | 0.97 |
| 3.53 | CB-76 CYS | 0.97 | 0.97 |
| 3.32 | CB-81 SER | 1 | 0.97 |
| 3.01 | CB-84 LEU | 1.14 | 0.98 |
| 5.37 | CB-85 SER | 1.16 | 0.98 |
| 3.08 | CB-87 ASP | 1.38 | 0.97 |
| 4.12 | CB-91 SER | 0.92 | 0.97 |

| | | | |
|---|---|---|---|
| 3.11 | CB-93 ASN | 1.18 | 0.98 |
| 3.85 | CB-94 CYS | 1.23 | 0.98 |
| 3.76 | CB-94 CYS | 1.11 | 0.97 |
| 5.27 | CG-105 MET | 0.98 | 0.97 |
| 3.98 | CB-105 MET | 1.05 | 0.97 |
| 3.18 | CB-105 MET | 0.87 | 0.97 |
| 3.86 | CB-108 TRP | 1.13 | 0.97 |
| 3.97 | CB-114 ARG | 1.08 | 0.97 |
| 3.92 | CB-115 CYS | 1 | 0.97 |
| 3.41 | CE-116 LYS | 1.13 | 0.97 |
| 3.04 | CD-116 LYS | 0.94 | 0.97 |
| 3.18 | CG1-124 ILE | 1.01 | 0.97 |
| 3.12 | CA-126 GLY | 0.92 | 0.97 |

**Extended Data Table 12.** Peak height of the difference peak in the Fc–Fo map for a candidate hydrogen on a methyl carbon ($CH_3$). The peak height is reported as the contour level (rmsd) at which the peak is no longer visible. The table also includes the atom label and residue of the binding atom, as well as the bond distance between the binding atom and the candidate hydrogen (Atom–H). Total observations: 25.

| **Peak rmsd** | **Binding atom** | **From Q peaks Atom-H (Å)** | **After refinement Atom-H (Å)** |
|---|---|---|---|
| 4.61 | CD1-8 LEU | 1.04 | 0.98 |
| 3.7 | CD1-8 LEU | 1.43 | 0.98 |
| 3.11 | CD1-8 LEU | 0.96 | 0.96 |
| 3.07 | CB-9 ALA | 0.99 | 0 occupancy |
| 3.86 | CE-12 MET | 1.16 | 0.98 |
| 3.25 | CE-12 MET | 1.11 | 0.98 |
| 5.03 | CD1-17 LEU | 0.91 | 0.97 |
| 3.24 | CD1-17 LEU | 1 | 0.97 |
| 4.64 | CD2-25 LEU | 0.91 | 0.97 |
| 3.9 | CD1-25 LEU | 1 | 0.97 |
| 3.58 | CD2-25 LEU | 0.93 | 0.97 |
| 3.55 | CG2-29 VAL | 0.94 | 0.98 |
| 3.74 | CB-31 ALA | 0.99 | 0.97 |
| 3.21 | CB-32 ALA | 1.14 | 0.98 |
| 3.96 | CG2-58 ILE | 1.26 | 0.97 |
| 3.35 | CD2-75 LEU | 0.81 | 0.97 |
| 3.05 | CD2-75 LEU | 1.03 | 0.97 |
| 3 | CD2-83 LEU | 0.87 | 0.97 |
| 4.16 | CG2-88 ILE | 1.11 | 0.99 |
| 3.77 | CG2-88 ILE | 1.01 | 0.97 |
| 3.49 | CG2-88 ILE | 1.05 | 0.97 |
| 3.18 | CB-95 ALA | 1.06 | 0.97 |
| 3.16 | CD1-98 ILE | 1.06 | 0.97 |
| 3.1 | CG2-98 ILE | 1.21 | 0.97 |
| 3.93 | CG2-99 VAL | 0.82 | 0.97 |
